# Multimodal data analysis reveals asynchronous aging dynamics across female reproductive organs

**DOI:** 10.1101/2025.05.16.654406

**Authors:** Oleksandra Soldatkina, Laura Ventura-San Pedro, Allal El Hommad, Jose Miguel Ramirez, Aida Ripoll-Cladellas, Maria Sopena-Rios, Jaume Ordi, Natàlia Pujol-Gualdo, Marta Melé

## Abstract

Female reproductive aging is a complex process with profound systemic health implications, yet the molecular and structural dynamics of aging across reproductive organs and tissues remain largely unexplored. Here, we integrate deep learning-based analysis of 1,112 histological images with RNA-seq data from 659 samples across seven reproductive organs in 304 female donors aged 20-70 years. We show that female reproductive organs and tissues have asynchronous aging dynamics: while the ovary and vagina age gradually, the uterus undergoes an abrupt transcriptional and cellular transition around the age of menopause. Tissue segmentation highlights that the myometrium in the uterine wall is the most age-affected tissue, marked by extracellular matrix remodeling and immune activation. Across reproductive organs, the epithelial tissue is also strongly affected by age, with the vaginal epithelium showing a unique sharp menopausal transition. Integration via multi-omics factor analysis links these tissue-specific histological transformations to specific molecular shifts, many with nonlinear expression trajectories and enriched in heritable reproductive traits such as pelvic organ prolapse and age at menarche. These findings position menopause as a key inflection point in female aging and provide insights with tissue-specific focus to support healthier menopausal transitions and reduce age-related disease risk.

## INTRODUCTION

Female reproductive aging is a complex biological process marked by the gradual decline in ovarian function and fertility, followed by menopause, the permanent cessation of menstruation^1^. This transition reflects the depletion of ovarian follicles and a corresponding drop in estrogen production, but its consequences extend far beyond fertility loss. Notably, the onset of menopause has been associated with an increased risk of many different chronic conditions^1,2^ such as diabetes^3^, cardiovascular disease^4,5^, osteoporosis^6^, and neurodegenerative disorders^7^, underscoring its systemic physiological impact. Despite its clinical impact, female reproductive aging is one of the most understudied topics in the aging field. As highlighted by Gilmer, G. et al.^8^, preclinical aging models rarely incorporate female aging trajectories, with less than 1% including menopausal phenotypes. Furthermore, most of the human research on reproductive decline has focused on the changes leading up to menopause, with far less attention paid to the biological transformations that occur during and after this transition^9^. Additionally, while large-scale genetic studies have uncovered loci associated with menopause timing^10^, the molecular mechanisms and tissue-specific manifestations of these signals remain poorly understood.

Recent computational tools using machine learning can resolve organ-level changes with high spatial and temporal precision^11,12^. In particular, histological image analysis has emerged as a powerful approach to uncover morphological aging patterns. Deep learning methods, such as convolutional neural networks (CNNs), have significantly advanced histopathology analysis by allowing the systematic identification of many phenotype-related morphological changes^13–15^, including aging-related changes across 26 human organs^16^ and the mouse kidney^17^. CNN-based aging clocks have been developed using skin biopsy images to estimate mortality risk^18^ or predict clinical outcomes from human hippocampal images^18,19^. More recently, Vision Transformers (ViTs), which utilize self-attention to capture complex spatial relationships^20,21^, have outperformed CNNs in some tasks^21–24^. ViTs-based approaches have revealed age-related changes in tissue-organ composition^20,21^ in breast and skin and further improved biological age prediction when integrated with gene expression data^25^. Thus, deep learning holds enormous potential to dissect aspects of human health including aging dynamics.

Currently, our biological understanding of reproductive aging — particularly across organs — remains limited and fragmented. Most studies have focused on studying the molecular changes during reproductive decline in specific organs, particularly the ovary^26–31^. These have highlighted core pathways such as DNA damage response and identified *FOXP1* as a key transcriptional regulator for ovarian senescence^24,25^. Beyond the ovary, smaller-scale efforts have identified expression changes with age in the fallopian tube^28^, endometrium^31^, and myometrium^29^, revealing key aging pathways, transcriptional regulators, and altered intercellular interactions. However, these studies are largely driven by interest in fertility and early reproductive decline, and are constrained by limited sample sizes and organ-specific scope. As a result, it remains unclear whether reproductive aging follows a coordinated molecular program across tissues or unfolds at different rates and inflection points^30^. Moreover, to date, only a single gene from murine data has been annotated as related to menopause in the Gene Ontology database — one of the most widely used tools for pathway analysis^32,33^. Together, this highlights a critical gap in the biological characterisation and functional annotation of the molecular processes underlying menopause across female reproductive organs.

Here we integrate 1,112 histological images paired with 659 RNA-sequencing (RNA-seq) samples from seven reproductive organs from the GTEx project^34^ to unravel the histological and molecular dynamics of female reproductive aging. Using machine learning on both histological and molecular data, we find that female reproductive tissues exhibit distinct aging trajectories within the same organ, suggesting that aging dynamics is asynchronous across reproductive tissues. For example, the ovary shows early, gradual changes while the uterus undergoes a sharp menopausal transition. In addition, we identify thousands of age-associated genes across organs, associated to extracellular matrix remodeling, immune response, oogenesis, and angiogenesis, and enriched in reproductive aging, hormonal and clinical traits. Our work unravels molecular and histological changes underlying female reproductive aging across organs which can help develop interventions that mitigate menopause-related decline and promote a healthier lifespan^35^.

## RESULTS

### Deep learning uncovers different aging dynamics across female reproductive organs

Deep learning provides a powerful approach for visualizing and quantifying age-related morphological changes in histology images^36,37^. To systematically investigate female reproductive aging, we analyzed 1,112 histological images from the GTEx v10 dataset, covering seven key female reproductive organs — uterus, ovary, vagina, breast, endocervix, ectocervix and fallopian tubes — from donors aged 20 to 70 years (Fig. 1A). First, we assessed whether deep learning models could reliably distinguish histological images from young or old donors. To address this, we fine-tuned a pretrained CNN (VGG-19) to classify histological images into young (<35 years) and old (>60 years) in the four organs with sufficient data (>10 samples per age group) — uterus, ovary, vagina, and breast, excluding middle-aged samples (35–60 years) (Fig. 1B, see Methods). In short, we extracted tiles of the same size for all sample images, and we get a probability of being classified as old per tile, that we can then average to get one value per sample. Validation accuracy exceeded 0.75 per tile and reached 1.0 per sample in the ovary, uterus, and vagina (Fig. 1C, Extended Data Table 1), indicating that CNN captures effectively the histological changes occurring across female reproductive organs through aging. In contrast, classification performance in the breast was lower (0.68 per tile, 0.79 per donor), likely driven by either more subtle or more heterogeneous age-related changes related to unannotated factors known to affect the breast histology^38–40,40,41^. Therefore, it was excluded from further image-based analyses in the current section.

**Fig.1:**
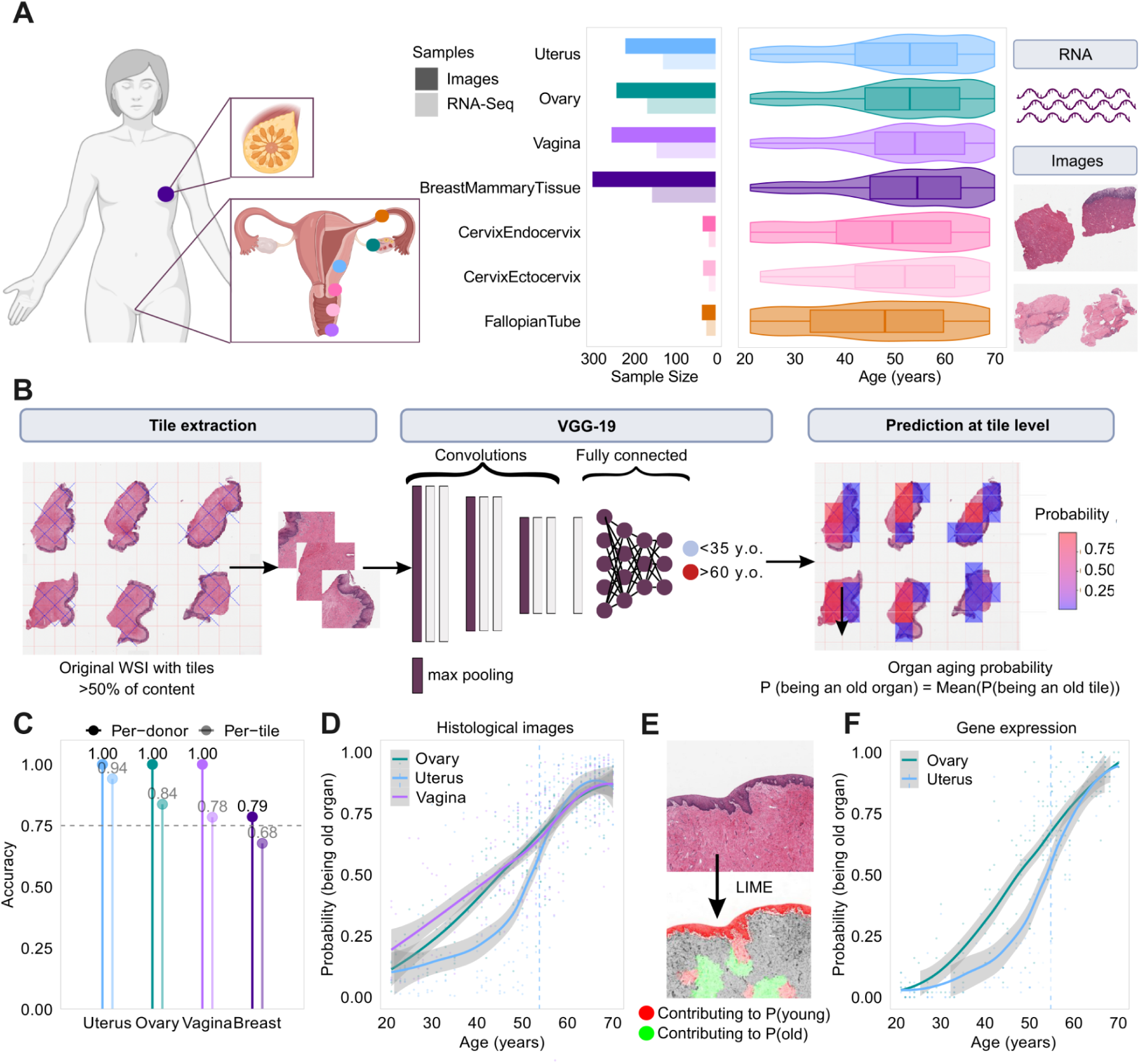
Histological and molecular aging trajectories across female reproductive organs are asynchronous. **A** Organs analyzed, sample sizes, and donor age distribution of image samples. **B** Deep learning workflow for predicting organ age group from histological images. **C** Validation accuracy across organs, evaluated per tile and per donor (tile-averaged). **D** Image classification-based aging trajectories for the uterus, ovary, and vagina; vertical dashed line marks the age of steepest change for nonlinear trajectory. **E** Vaginal histological image (top) with LIME interpretation map (bottom); red and green regions indicate areas that decrease or increase CNN model confidence, respectively. **F** Gene expression classification-based aging trajectories for the uterus and ovary; vertical dashed line marks the age of steepest change for nonlinear trajectory (only organs presenting classification accuracies >0.75 are shown).

Next, we wanted to test if reproductive organs have different aging trajectories. To do this, we applied the trained model to all samples to estimate the probability of being classified as ‘old’ (Fig. 1D) and used these probabilities across ages to infer organ-specific aging trajectories. In both ovary and vagina, the probability of being old increases linearly with chronological age (Davies test p.value>0.05, Extended Data Table 2). In contrast, the uterus has a nonlinear trajectory (Davies test p.value=3.48e-06), with most samples from 20 to 40 classified as young followed by a sudden transition to higher probability values starting around the age of 45 (Fig. 1D). Importantly, the steepest change in the uterus probabilities occurs at the age of 53 which is very close to the average age of natural menopause at 51 years^42^. Consistent with this, ovarian samples have significantly higher probabilities of being old than uterus images from the same donor (exact binomial test p.value=6.75e-10) (Extended Data Fig. 1A). To address if specific histological features could drive the model predictions, we applied LIME (Local Interpretable Model-Agnostic Explanations)^43^ — an explainability method that uses image perturbations to identify areas of an image that are most informative for the CNN prediction. In the ovary and uterus, informative regions are distributed throughout the images (Extended Data Fig. 1B-C), suggesting that no specific histological structure is driving the age predictions. Conversely, in the vagina, predictions are predominantly influenced by the epithelium, consistent with previous observations of a vaginal epithelium thinning after menopause^44,45^ (Fig. 1E, Extended Data Fig. 1D).

To assess whether transcriptomic changes reflect the observed histological aging dynamics, we trained machine learning classifiers on RNA-seq data available from 659 matched samples (samples with both histological and RNA-seq data) (Fig. 1A) to distinguish young (<35 years old) from old (>60 years old) organs. Classification accuracy exceeded 0.75 in the ovary and uterus (Extended Data Fig. 1E, Extended Data Table 3), suggesting stronger transcriptomic aging signals in these organs. Inferred aging trajectories from gene expression (Fig. 1F) show gradual changes in the ovary with age (Davies test, p.value=0.305), whereas the uterus exhibits an abrupt shift at age 51 (Davies test, p.value= 4.31e-06), similar to the histological predictions (Fig. 1E). We then used a sliding-window approach to identify nonlinear changes, by comparing gene expression between adjacent 10-year intervals within a 20-year window (see Methods) and shifting the center by 1 year. Consistently, we identified the highest number of age-differentially expressed genes (DEGs) — 189 DEGs — in the uterus around age 50 (Extended Data Fig. 1F). In contrast, we detected fewer age-related DEGs in the ovary and vagina, consistent with more progressive transcriptional changes that are harder to detect in shorter age intervals. Overall, our results show that histological and transcriptional aging dynamics are asynchronous across female reproductive organs with the ovary and vagina starting to age gradually earlier in life and uterus undergoing an abrupt change around the age of menopause.

### Myometrium and epithelial tissues are primary sites of reproductive tissue aging

Organs are composed of distinct tissues organized in complex structures that may be differentially impacted by aging. To study different tissues (e.g., epithelium, adipose) and tissue structures (e.g., vessels) within each organ, we applied a segmentation method based on the vision transformer Dino ViT-S/8^46^ pretrained on histological images of 23 healthy non-reproductive organs (except for breast)^21^. We first fine-tuned the model on whole-slide images from reproductive organs, including the uterus, ovary, vagina, fallopian tubes, endocervix, and ectocervix, ensuring that it could accurately represent their histological features. This fine-tuning produced 384 ViT-derived features per tile (Fig. 2A), which capture semantic histological information such as tissue architecture and cellular morphology^21^. We clustered the tile-level representations within each tissue structure using a limited set of structure-annotated tiles (Fig. 2A). Clustering accuracy exceeds 97% for all organs (Extended Data Fig. 2A). As expected, when combining all tiles from different organs together, the tiles cluster by tissue and tissue structure rather than by organ or donor (Fig. 2B), suggesting that ViT-derived features robustly represent functional tissue architecture across reproductive organs (Extended Data Fig. 2B-H). Next, for each tissue and tissue structure, we quantified the proportion of variation in histological image features attributable to age, while controlling for other demographic variables and batch effects (Fig. 2C, Methods). The median variance explained by age ranged from 2 to 9% across tissues, with the myometrium showing the largest median (9%). Interestingly, some myometrium features have over 40% of their variation explained by age (Fig. 2C). Visual inspection of myometrium tiles across values of the most age-variable feature revealed pronounced differences between young and old donors, with lighter pink areas in the old donors (Fig. 2C, Extended Data Fig. 2I, see Methods). Comparatively, the endometrium has a lower proportion of variation explained by age, even after accounting for its lower representation in the image tiles (see Methods). This is somewhat unexpected, given that endometrium transitions to a non-cycling atrophic state during menopause. However, this lower age-associated variation may reflect the dominant influence of other factors —such as the regular cyclical remodeling that occurs during the menstrual cycle prior to menopause or the use of hormonal replacement therapy afterwards. In the vagina, endocervix, ectocervix, and fallopian tubes, the epithelium is the most affected tissue, which expands on previously reported morphological and physiological menopause-related changes to vaginal and cervical epithelial tissue^45,47^. Visual inspection of epithelium tiles across values of the most age-associated feature in vagina showed increased nuclear-to-cytoplasm ratio in old donors (Fig. 2C, Extended Data Fig. 2J), notably with the same feature changing with age in the epithelia of the other reproductive organs (Extended Data Fig. 2K-M). In the breast, lobules — structures composed of an epithelial bilayer responsible for milk production during lactation — have the highest age-explained variation (Fig. 2C). Across reproductive organs, the most age-variable tissue features reveal distinct aging dynamics (Extended Data Fig 3A-D) — linear in the tissues of the ovary, but a mix of linear and nonlinear trajectories between the tissues of the uterus, vagina and breast.

**Fig. 2:**
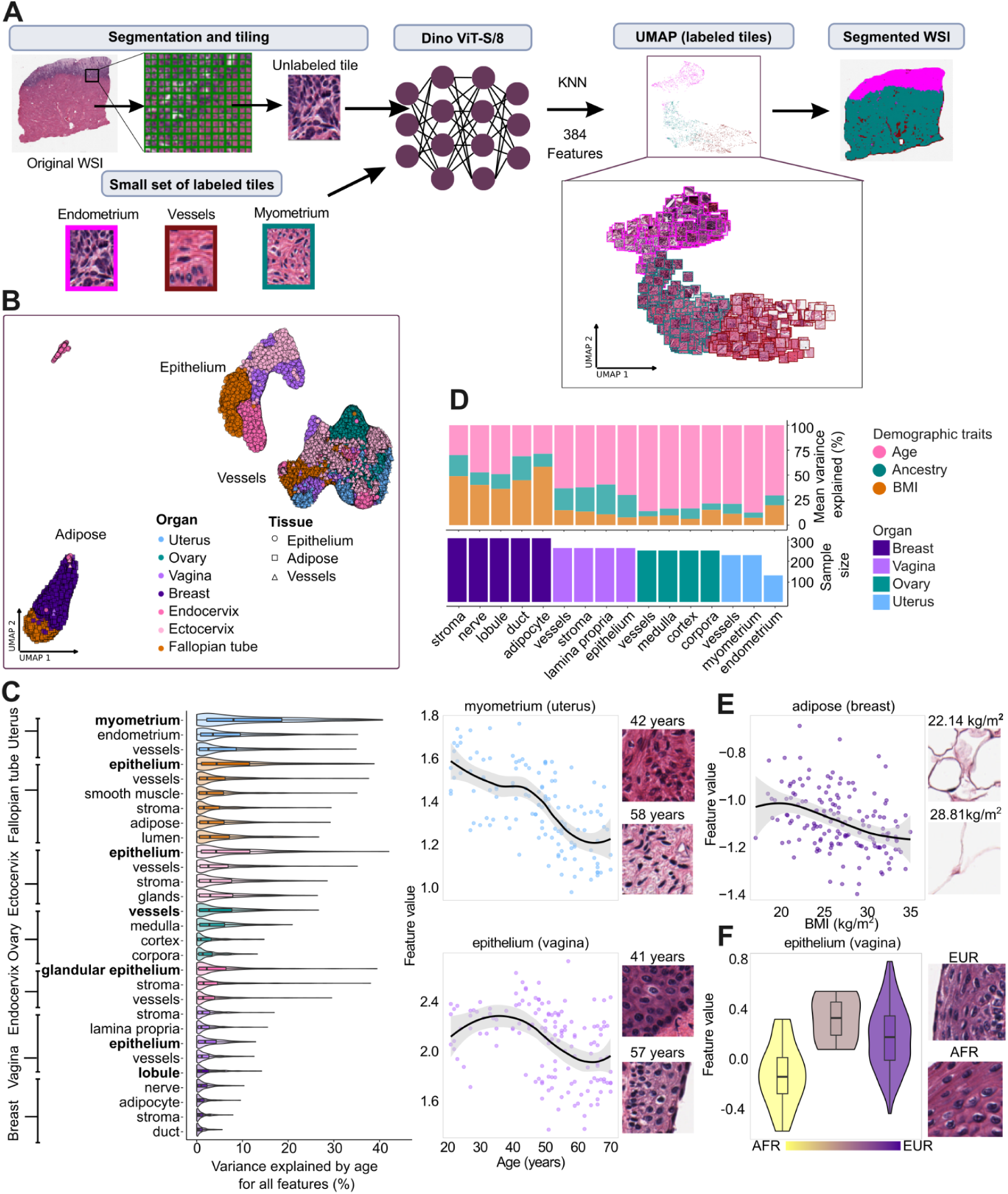
Vision transformer–based segmentation reveals age as a primary driver of histological variability across female reproductive tissues. **A** ViT-based segmentation workflow for whole slide image segmentation into tissue and tissue structure tiles. **B** UMAP projections of a subset of identified tissues and tissue structures in reproductive organs. **C** Variance explained by age for each tissue and tissue structure with illustrative examples of feature aging trajectories and representative tiles for myometrium (top right) and vaginal epithelium (bottom right). **D** Mean proportion of variance explained by age, genetic ancestry and BMI across tissues. **E** Relationship between a BMI-variable image feature and BMI for adipose tissue (left) with representative tile images (right) **F** Relationship between an ancestry-variable image feature and genetic ancestry for vaginal epithelium (left) with representative tile images (right).

We next compared the impact of age, body mass index (BMI) and genetic ancestry on histological variation across tissues (Fig. 2D, Extended Data Fig. 3E). Age accounted for the largest proportion of variance in almost all tissues. BMI had its strongest effect in the breast, particularly in adipocytes, where it explained up to 2.6% of variance (Fig. 2D,E, Extended Data Fig. 3F-G), likely reflecting detectable histological changes in adipocytes with BMI. In contrast, genetic ancestry explained more variance than BMI in all four vaginal tissues. These differences cannot be attributed to unequal ancestry distribution across organs (Extended Data Fig. 3E). Visual inspection of ancestry-informative features reveals denser squamous epithelial cells in African ancestry donors compared to European donors of the same age (Fig. 2F, Extended Data Fig. 3H), suggesting that vaginal histology may vary across ancestries.

Overall, our histological segmented feature image analysis shows that aging affects most tissues and tissue structures of the female reproductive tract, with myometrium being the most affected and epithelial tissues having a coordinated response to aging across all reproductive organs.

### Menopause drives divergent aging trajectories within reproductive tissues and across organs

After identifying distinct aging trajectories at the organ level (Fig. 1D), we next investigated aging rates at higher granularity to assess whether reproductive tissues age at different paces within the same organ. To this end, we trained elastic net classifiers on tile-level image features to distinguish young (<35 years old) from old (>60 years old) samples (see Methods). We focused on tissues with at least 10 samples per age group, and only kept the highest performing classifiers per tissue with a minimum accuracy ≥0.75 (myometrium and vessels in uterus, epithelium, lamina propria and stroma in vagina, and corpora, cortex and vessels in ovary, Extended Data Fig. 4A). We applied these classifiers to predict histological age on tiles from all donors, including those aged 35–60 years, enabling us to trace intermediate histological trajectories based on per-sample predictions. In the uterus, both myometrium and vessels exhibited synchronous nonlinear aging trajectories with sharp transition around the age of natural menopause (51 years) (Davies test, p.value < 0.05, Fig. 3A, Extended Data Table 2), indicating a coordinated tissue-wide response during menopause. In contrast, vaginal tissues had heterogeneous aging dynamics (Fig. 3B): the lamina propria and stroma aged gradually (Davies test, p.value > 0.05), whereas the epithelium followed a menopause-associated nonlinear trajectory (Davies test, p.value = 0.027), with the steepest change at age 51.7 years. Conversely, ovarian tissue trajectories are highly similar though the ovarian cortex displays a subtle nonlinear trend (Davies test, p.value=0.04669), suggesting only mild differences in aging dynamics across ovarian tissues.

**Fig. 3.**
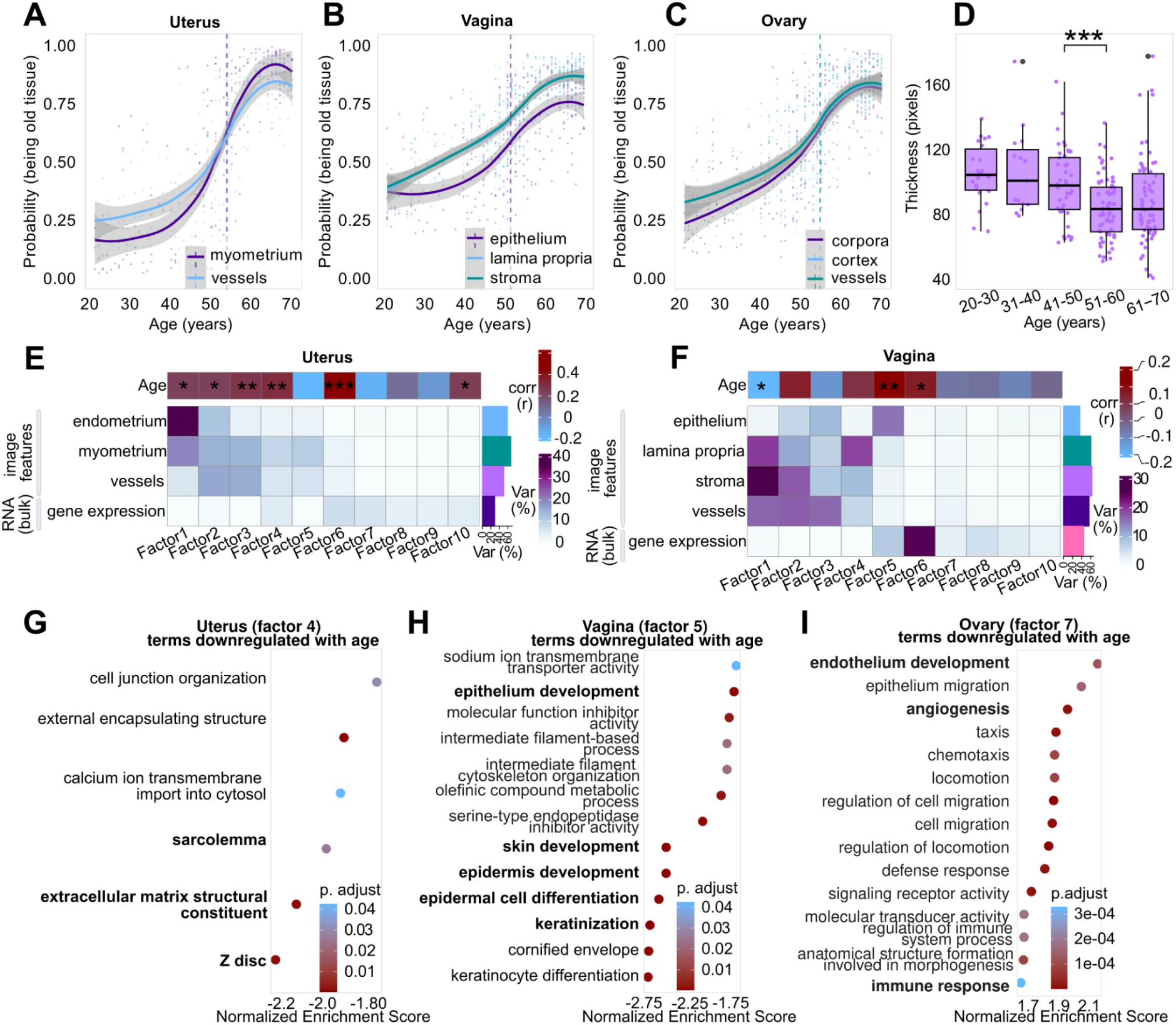
Multimodal analysis shows within-organ tissue-specific aging dynamics reflecting distinct susceptibility to menopause. **A-C** Predicted aging trajectories for uterus (A), vagina (B), and ovary (C) based on ViT-derived features, with dashed lines marking the most rapid transition. **D** Vaginal epithelial thickness with age; *** indicates FDR<0.0005 (Wilcox pairwise test). **E-F** Multi-Omics Factor Analysis (MOFA) results for uterus (E) and vagina (F): heatmaps show the variance explained by 10 latent factors across modalities (bottom) and their correlation with donor age (top). Asterisks denote significance (*FDR<0.05 & >0.005, **FDR<0.005 & >0.0005, ***FDR<0.0005). **G-I** Gene set enrichment analysis (GSEA) using Gene Ontology (set size >40 genes) for genes negatively contributing to Factor 4 in the uterus (G), Factor 5 in the vagina (H), and genes positively contributing to Factor 7 in the ovary (I).

Next, we assessed differences in tissue proportions during aging. We observed that several tissues change proportions with age (Extended Data Fig. 4B), but we could not identify nonlinear aging dynamics across tissues (Davies test p-value>0.05, Extended Data Table 17) likely because sampling differences add a significant amount of noise to tissue proportion estimates across age groups (Extended Data Fig. 4B-D). To validate whether a sharp remodeling event occurs in the vaginal epithelium, we directly measured epithelium thickness from the segmented histological images using CellProfiler^49^ and observed a significant thinning specifically between 40’s and 50’s decades (Fig. 3D, Wilcoxon test, p-adj = 0.006), consistent with a menopause-associated transition that is not captured by proportion estimates, which shows a monotonic proportion decay with age (Davies test, p.value= 0.791, Extended Data Fig. 4B). This discrepancy likely reflects the limitations of proportion-based estimates, which are highly sensitive to how tissues are sampled and sectioned across individuals (Extended data Fig. 4B-D). Similarly, trajectory inference on cell type deconvolution estimates from gene expression reveal only a few age-associated shifts in cell type composition, suggesting that cell type proportion estimates from gene expression may be too noisy to perform reliable trajectory inference (Extended Data Fig. 4E-G). Thus, histological segmented feature image analysis is a more robust tool for detecting age-related tissue reorganization, as it appears to be less influenced by sampling variability compared to tissue or cell type proportion estimates.

**Fig 4.**
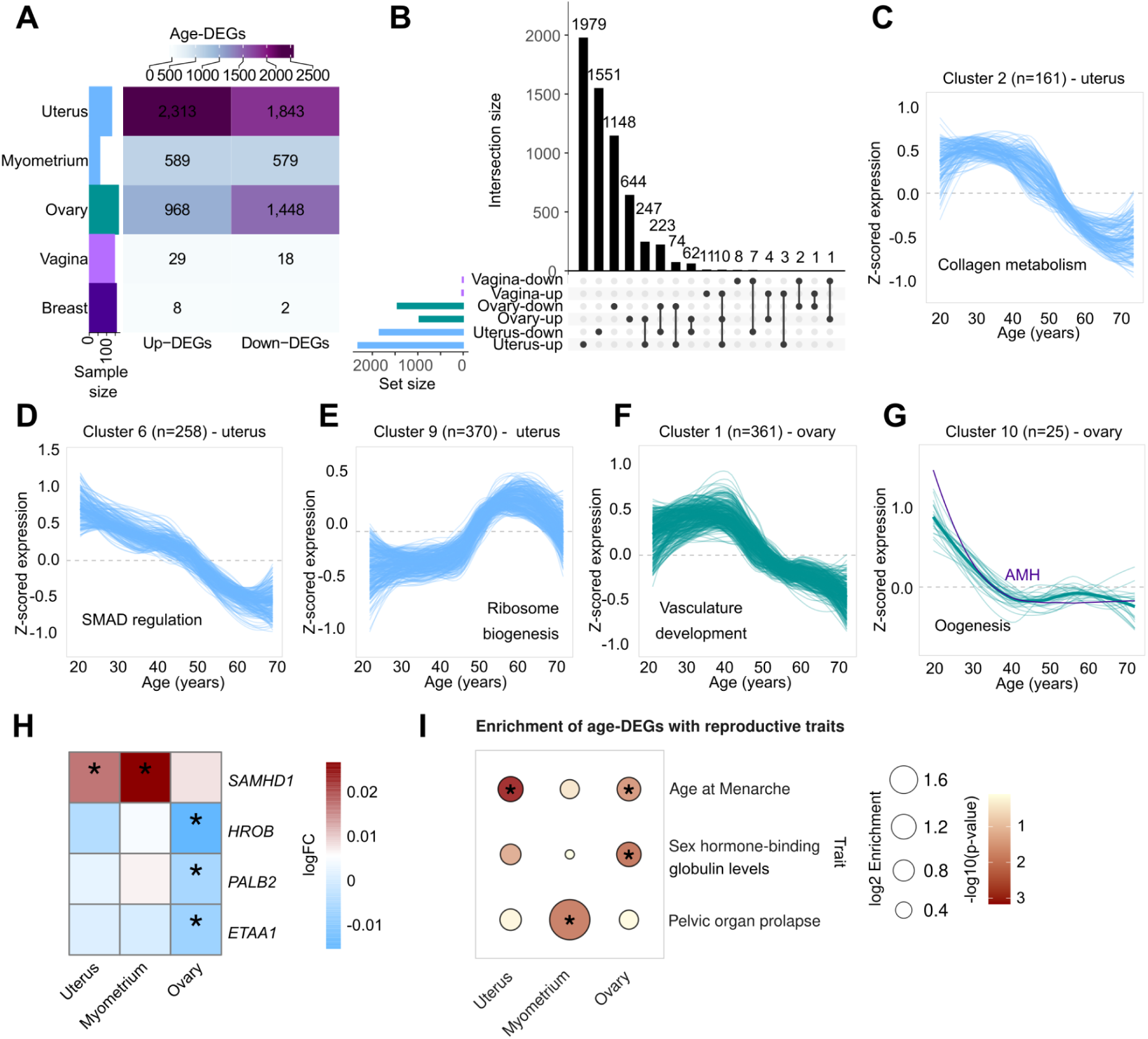
Nonlinear expression dynamics and trait-linked gene enrichment during reproductive aging. **A** Number of age-differentially expressed genes (age-DEGs) identified per organ or in myometrium-only samples. **B** Shared upregulated and downregulated age-DEGs across the uterus, ovary, and vagina. **C-E** Uterine gene trajectory clusters enriched in (C) collagen metabolism, (D) SMAD regulation, and (E) ribosome biogenesis. **F - G** Ovarian gene trajectory clusters enriched in (F) vascular development and (G) oogenesis. **H** Heatmap of age-DEGs overlapping with large-effect genes for age at natural menopause. Asterisks indicate significant age-DEGs (FDR<0.05). **I** Enrichment of age-DEGs with reproductive traits. Asterisks indicate significant enrichments (Fisher test; FDR<0.05).

Our vision transformer–based approach reveals tissue-specific aging trajectories, enabling detection of sharp menopausal transitions that are missed by traditional bulk or proportion-based methods.

### Multi-modal factor analysis uncovers molecular drivers of tissue-specific reproductive aging

Next, we wanted to identify transcriptomic changes associated with the observed tissue-specific histological changes with age. However, a key challenge is that available GTEx expression data is derived from bulk RNA-sequencing, capturing mixed signals from multiple tissue types within each organ. To address this, we applied Multi-Omics Factor Analysis (MOFA)^49,50^ to integrate ViT-derived histological features per organ-associated tissue with gene expression data per organ (see Methods). MOFA extracts latent factors: low-dimensional variables that capture shared and specific sources of variation across data modalities. As data modalities for each organ, we used 384 ViT-derived image features per tissue and tissue structure averaged per donor and gene expression profiles from the 5,000 most variable genes.

Latent factors explained a greater proportion of variance in image features than in gene expression (Fig. 3E-F, Extended Data Fig. 5A-B, Extended Data Table 4). The breast stroma histology had the largest variance explained (77%, Extended Data Fig. 5B), suggesting that unmeasured biological or technical factors may underlie its lower classification performance (Fig. 1C). Most factors explain variation in a single modality — either histology (e.g., Factors 1-3 in the uterus (Fig. 3E)) or gene expression (Factors 6-10 in the uterus (Fig. 3E)). However, a subset of factors capture shared variation across both modalities and are significantly associated with age. In the uterus, Factors 4 and 6 captured age-associated changes (r=0.31 and p-adj=8.072e-04, r=0.47 and p-adj=1.284e-07) in myometrium and vessel histology with gene expression. Genes whose expression is down-regulated the most with Factor 4 are associated with pathways related to extracellular matrix and muscle cell structure and function (Fig. 3G, Extended Data Table 5). The enrichment in the extracellular matrix likely reflects the visible histological differences between young and old myometrial images, in which the light pink areas predominant in old donors correspond to collagen deposition (Fig. 2D, Extended Data Fig. 5C)^30,51^. In contrast, Factor 4 up-regulated genes are enriched in chromatin remodelling (Extended Data Fig. 5D), likely due to the epigenetic reprogramming that accompanies hormonal withdrawal and tissue remodeling during the menopausal transition^52,53^. In the vagina, Factor 5 captures coordinated aging (r=0.23 and p-adj=0.005) in epithelium histology with gene expression changes mostly involved in epithelium development (Fig. 3H). In the ovary, Factor 7 connected aging-related shifts (r=-0.46 and p-adj=3.397e-10) across the cortex, vessels, corpora albicantia, and gene expression changes associated with angiogenesis pathway (Fig 3I, Extended Data Fig. 5G). Angiogenesis is known to undergo age-related suppression^26,54^. Finally, two factors — Factor 5 in vaginal epithelium and Factor 6 in uterine myometrium — are associated with up-regulation of immune system-related terms (Extended Data Fig. 5E-F), consistent with a general increase in inflammation during aging^44,51^.

By integrating histological and gene expression data, we identify key molecular pathways underlying age-related tissue-specific changes, providing a direct link between histological transformations and coordinated gene expression shifts.

### Gene expression shifts drive functional reproductive decline and associate with reproductive traits

Gene expression changes with age can follow gradual or nonlinear trajectories, with some genes shifting progressively and others exhibiting sharp transitions around menopause. To resolve these temporal dynamics, we performed differential expression analysis on each reproductive organ, controlling for tissue composition (see Methods). Since not all the samples have endometrium, we also performed differential gene expression analysis in the uterus samples only composed of myometrium separately (see Methods). The uterus and ovary had the largest number of age differentially expressed genes (age-DEGs) (Fig. 4A, Extended Data Tables 7,9). Among these, several age-DEGs are shared across organs (Fig. 4B), with the largest overlap found between genes upregulated in both ovary and uterus followed by those downregulated in both organs. From the up and downregulated myometrium age-DEGs, there is an overlap of 82 and 87% with uterus up and downregulated age-DEGs, respectively.

We next clustered age-DEGs based on their expression trajectories and identified 14 clusters in uterus, 10 in ovary and 2 in vagina, proportional to the number of age-DEGs per organ (Extended Data Fig. 6A-B, 7A). Functional enrichment analysis assigned 11 of these clusters into distinct functional categories (Extended Data Tables 8, 10, 11). In the uterus, six clusters were functionally annotated, with three exhibiting abrupt expression changes around menopause (Fig. 4C-E). Two of these clusters show a sharp decline in gene expression. One is enriched in extracellular matrix organization and collagen metabolism (Fig. 4C) — likely reflecting histological remodeling of the myometrium during menopause (Fig. 2D). The second cluster (Fig. 4D), also downregulated at menopause, is involved in SMAD protein signal transduction within the Transforming Growth Factor beta (TGF-β) pathway, previously implicated in endometrial dysfunction^55,56^. Notably, TGF-β pathway activity is tightly regulated throughout the menstrual cycle, suggesting that its downregulation may reflect disrupted stromal-epithelial signaling during the menopausal transition^57^. In contrast, the third and largest cluster shows a rapid, synchronous increase in gene expression at menopause (Fig. 4E), primarily related to ribosome biogenesis and rRNA processing. The postmenopausal upregulation of ribosomal genes may reflect increased translational demands associated with senescence, senescence-associated secretory phenotype (SASP) production, or tissue remodeling in response to the chronic inflammation^58,59^.

In the ovary, all four functionally enriched gene clusters show nonlinear expression dynamics (Extended Data Figs. 7A). One cluster has an abrupt trajectory change at the age of natural menopause and it is associated with decrease of vasculature development genes, consistent with the age-related angiogenesis suppression in the ovary^26,60^ (Fig. 4F). Another cluster shows much earlier downregulation and is enriched in female reproduction and oogenesis genes including *AMH,* used as biomarker of ovarian reserve^61^ and consistent with ovarian fertility decay starting earlier in life.

To place these molecular changes in the context of reproductive lifespan, we next examined two key reproductive aging traits: age at menarche and age at natural menopause. Both traits are influenced by hundreds of common genetic variants and nearby genes, identified by large-scale genome-wide association studies (GWAS)^62,63^. In total, we find that age-DEGs in the uterus, ovary, and/or the myometrium overlap with 65 menopause associated GWAS genes (Extended Data Fig. 7B) although this overlap is not higher than expected by chance (45 genes uterus-specific, Fisher test, OR= 0.82 [0.57-1.14], p.value=0.26). Notably, four of the nine genes reported in analyses of rare protein-coding variants strongly impacting age at natural menopause are age-DEGs (*SAMHD1*, *HROB*, *PALB1*, and *ETAA1*)^64^*. SAMHD1* is upregulated in the uterus and myometrium-only samples, while *HROB*, *PALB1* and *ETAA1* are downregulated in the aging ovary (Fig. 4H). We find significant enrichment of age-DEGs in genes associated with age at menarche in the uterus (161 genes, Fisher test, OR = 1.39 [1.14-1.68], p.adj=0.002) and to a lesser extent, in the ovary (94 genes, Fisher test, OR = 1.32 [1.04-1.66], p.adj=0.03) (Fig. 4I, Extended Data Table 14). We also tested whether our age-DEGs significantly overlapped with genes associated with other female reproductive traits including clinical outcomes and hormonal traits identified in ten different GWAS studies (Extended Data Table 13). Among clinical traits, uterine myometrium age-DEGs are enriched for genes associated with pelvic organ prolapse (8 genes, OR= 3.16 [1.26-6.94], p-adj=0.02). In the ovary, age-DEGs overlapped with GWAS genes for sex hormone-binding globulin levels (107 genes, OR = 1.36 [1.09-1.69], p-adj=0.005). These findings highlight the potential tissue-specific roles for genes influencing reproductive aging traits such as age at menarche and age at menopause, and reinforce that its impact extends beyond the ovary and ovarian function, highlighting the importance of uterine and myometrial dynamics – alongside ovarian decline – in shaping female reproductive aging.

## DISCUSSION

In this study, we present the first large-scale, tissue-resolved analysis of female reproductive aging across seven human organs. By integrating deep learning-based histological profiling with transcriptomic data from 659 donors, we reconstruct molecular and morphological aging trajectories of the uterus, ovary, vagina, cervix, fallopian tubes, and breast. Our approach leverages convolutional and vision transformer models to achieve tissue-level resolution, enabling the discovery of asynchronous aging dynamics within and across reproductive organs. This work fills a major gap in aging research, where reproductive organs —particularly beyond the ovary— remain understudied despite menopause’s strong association with increased risk of systemic diseases^1–7^. By generating an integrated tissue atlas of human female reproductive aging, our study establishes a foundation for understanding how organ-specific aging patterns contribute to health outcomes across the menopausal transition.

One of the most striking patterns resulting from this work is that menopause marks a biological inflection point that triggers coordinated, rapid remodeling across multiple reproductive tissues. This supports the emerging view of aging as a nonlinear process marked by periods of accelerated change^65–67^. Such dynamic transition may help explain the sharp increase in chronic disease risk observed in females after menopause^1,68^. Furthermore, recent studies using blood-derived biomarkers suggest that aging can be organ-specific and influence diseases affecting the specific organs^69,70^, however these inferences often lack direct organ-level validation. Here, we provide such validation by showing that human female reproductive organs age in distinct and asynchronous ways. The ovary and vagina show early, gradual aging, consistent with the well-established decline in fertility beginning a decade before menopause^71^ (Fig. 1D). In contrast, the uterus undergoes abrupt remodeling around the age of menopause (Fig. 1D). Within these organs, the myometrium and the epithelium, respectively, show the most pronounced transitions (Fig. 3A-B), highlighting that aging is not only organ-specific but also asynchronous across tissues within the same organ. Our results emphasize the need to understand the dynamics, timing and tissue-specificity of aging, critical for identifying deviations from normative reproductive aging trajectories, which could inform individualized risk stratification and preventive strategies.

At the molecular level, we find that the aging myometrium is marked by increased collagen deposition and downregulation of muscle-related pathways (Fig. 3G, Fig. 4C). These changes are accompanied by enrichment of age-associated genes at pelvic organ prolapse GWAS loci^72^ (Fig. 4I), suggesting a mechanistic link between menopause-associated tissue weakening and this highly prevalent condition, which is a major cause of hysterectomy^73,74^ (surgical removal of the uterus). Our findings position collagen remodeling as a hallmark of myometrial aging, and raise the possibility that circulating markers of extracellular matrix turnover — such as procollagen peptides and matrix metalloproteinases — could serve as non-invasive biomarkers of uterine tissue health, as already explored in cardiovascular^75^ and liver diseases^76^. These markers may also aid in early risk assessment for pelvic organ prolapse and pregnancy complications, given known association between myometrial dysfunction and adverse obstetric outcomes, including implantation failure, preterm birth, labor dystocia, and uterine atony^77–79^, complications that are becoming more prevalent with rising maternal age^80^. In parallel, we observe coordinated epithelial remodeling across reproductive tissues. While vaginal epithelial thinning is a well-known hallmark of menopause^45,47^, our analysis reveals broader changes in epithelial organization — including in the cervix and fallopian tubes —, such as increased nuclear-to-cytoplasm ratio (Fig. 2C, Extended Data Fig. 2J), that occur independently of thickness. The epithelial aging may compromise barrier integrity and innate immunity^81,82^, potentially contributing to pelvic inflammatory disease later in life^81^. These findings deepen our mechanistic understanding of the vaginal signs and symptoms characteristic of genitourinary syndrome of menopause — a highly prevalent but often underdiagnosed post-menopausal syndrome with a genital affectation characterised by vaginal dryness, pain and related atrophic changes^83^ — further extending its implications into the upper reproductive tract.

Furthermore, we offer a framework to explore inter-individual variability in both intra- and inter-organ asynchrony in female reproductive aging, which may underlie differences in menopause timing and its associated disease risks, including type 2 diabetes, osteoporosis, and hormone-sensitive cancers^63^. While late menopause is linked to increased longevity^84^ and higher risk of hormonally-driven cancers^63,64^, it remains unclear whether early menopause reflects accelerated systemic aging. Organ-specific variability in aging trajectories may also underlie the clinical heterogeneity of gynecological conditions such as polycystic ovary syndrome and uterine fibroids^85,86^. Additionally, our enrichment analysis shows that uterine and ovarian age-associated genes (e.g., *SH2B1*, *THRB*, *INHBB*, *ERBB3*) (Fig. 4H) are over-represented in loci associated with menarche timing, providing functional, tissue-specific context for signals from large-scale GWAS and supporting the idea that variability in early developmental events may influence later-life reproductive aging. Although enrichment at menopause loci was weaker — likely due to smaller discovery sample sizes (∼200,000 women versus ∼800,000 for menarche) — these findings suggest that genes influencing early reproductive milestones may also shape later-life tissue aging trajectories.

Despite these advances, our study has some limitations. The GTEx dataset lacks detailed reproductive metadata, including menopausal status, parity, breastfeeding history, hormone therapy use, and menstrual cycle phase. Indeed, while we expected stronger aging signatures in breast, the observed patterns were relatively muted, possibly due to confounding factors such as parity^38–40^, lactation history^40,41^, or highly heterogeneous histological sampling. Moreover, our findings suggest a significant role of ancestry in shaping reproductive tissue architecture (Fig. 2D-F), underscoring the importance of population diversity in aging research. In parallel, our analysis highlights candidate genes that warrant functional follow-up. One such example is *SAMHD1*, a top menopause-associated gene^64^ that increases expression with age in the uterus. While prior studies have linked *SAMHD1* deleterious variants to delayed menopause and proposed its inhibition to enhance ovarian response in IVF^64^, its effects on uterine biology remain unknown. Further work is needed to assess whether modulating *SAMHD1* influences uterine function or tissue remodeling, potentially informing future therapeutic strategies. To advance the field, future studies should leverage well-phenotyped longitudinal cohorts spanning broader age ranges and include ancestrally diverse populations. Expanding this multi-modal framework to additional organs and tissues, and incorporating single-cell and spatial transcriptomic approaches, will be key to uncovering cell-type–specific mechanisms driving female reproductive aging.

Our findings reposition menopause not merely as the end of the reproductive lifespan, but as a biological inflection point that initiates divergent aging trajectories across reproductive tissues. By linking these tissue-specific transitions to clinical conditions such as pelvic organ prolapse, our work provides mechanistic insight into the origins of common postmenopausal disorders. More broadly, this study demonstrates the power of a tissue-resolved, organ-level approach to study female reproductive aging — one that can inform precision strategies to monitor, predict, and ultimately mitigate age-related disease risk in women.

## METHODS

### GTEx data

The GTEx version 10 dataset consists of 19788 RNA-seq samples from 946 post-mortem donors and 54 tissues, with genotype data for 946 donors from whole genome sequencing available in a phased analysis freeze. The GTEx biospecimen collection, molecular phenotype data production, and quality control are described in detail in the GTEx v8 main paper^34^. GTEx v10 includes ∼12% more RNA-seq samples than v8, but these additional samples come from a subset of donors already present in v8, rather than new donors. GTEx v8 remains the primary source for whole-slide images (N=25,713) and genomes, covering all currently included donors. In this study, we analyze RNA-seq data from v10 alongside genomes and histological images from v8, focusing on female reproductive organs (uterus, ovary, vagina, breast, cervix and fallopian tube).

### Menopause timing definition

Given that most large hormonal shifts occur in the window of +/-5 years from last menstruation^87^, and the average age of menopause is 51^42^, we define the observed changes as menopause-timed if they peak in the age range of 45-55 years.

### Age ranges definition for classification

To capture biologically meaningful aging stages, we defined three age groups based on two reproductive milestones, ovarian aging onset and menopause onset. Ovarian aging typically begins around age 37^88^, while menopause timing usually occurs between 45 and 55^10^. Accordingly, we classified samples as young (20–35 years), middle-aged (36–59 years), and old (60–70 years) to reflect early ovarian decline and the menopausal transition (Extended Data Table 2).

### Image Filtering

Hematoxylin and eosin (H&E)-stained whole slide images (WSIs) were obtained from the GTEx Consortium in Aperio (.svs) format. We excluded misannotated whole slide images from the analyses based on histological annotations from GTEx. This included cases such as uterus samples annotated as vagina, organ samples containing a mixture of other organ slices, or samples with an unknown origin (exact terms of exclusion in Extended Data Table 15).

### Validation sets for image classifiers (CNNs and Elastic Nets)

We evaluated the predictive ability for young versus old classification using two different models (CNNs and Elastic nets) across histological images. Before training the predictive models, we designated a separate validation set to ensure unbiased performance evaluation. This validation set consists of 15 donors — 10 from old group (>60) and 5 from young (<35), to maintain the same ratio old/young as in the whole dataset — kept independent from the training and test sets. Additionally, to ensure consistency, we excluded from the models’ training 5 young donors (age <35) with one of the reproductive organs annotated as atrophic (Extended Data Table 15).

### Transfer Learning on Convolutional Neural Networks (CNNs) for Age Classification

To classify histological tiles as young or old, we trained separate CNN-based classifiers for each reproductive organ with sufficient samples (>10 per age group): breast, vagina, ovary, and uterus.

#### Image preprocessing

First we performed tissue segmentation on whole slide images using PyHIST^89^ and applying *Otsu* thresholding^90^ to separate organ content from the background. Non-overlapping tiles with >50% tissue content were retained for the uterus^91^, ovary, and vagina. In breast mammary tissue, a high proportion of transparent adipose content often confounded background detection, leading to a reduced number of extracted tiles.

To optimize tissue selection, we tested tissue content thresholds of 10%, 20%, 25%, and 50% using *Otsu* and graph-based segmentation. A 25% threshold with *Otsu* segmentation was selected as the optimal balance between tissue inclusion and background minimization. Tile sizes were set to 512×512 pixels for organs with large tissue slices (ovary, uterus, and breast) and 256×256 pixels for organs with smaller slices (vagina).

A total of 222, 245, 257, and 304 WSIs were analyzed, yielding 5,960, 6,000, 33,274, and 5,418 tiles for the uterus, ovary, vagina, and breast, respectively.

### Model architecture and training

We used the VGG-19 model^91^ pre-trained on the ImageNet dataset^91,92^. VGG-19 consists of 19 convolutional layers, each followed by a max-pooling layer that downsamples and aggregates information. Following the transfer learning paradigm, we froze the pre-trained VGG-19 weights and trained an additional fully connected layer with 1,024 nodes for binary classification (young vs. old).

We excluded middle-aged samples and the validation set, training only on young and old donors not present in these sets. The dataset was split into 75% training and 25% testing. To address class imbalance (fewer young samples), we applied WeightedRandomSampler(), assigning an inverse probability weighting to ensure equal representation of classes during training.

#### Hyperparameter tuning and optimization

The model was trained with a learning rate of 0.001, incorporating early stopping^93^ to prevent overfitting. Training stopped if validation accuracy did not improve for 10 consecutive epochs (patience parameter) or if the change in validation accuracy was below δ = 0.001.

To optimize model performance, we trained each model with different dropout and gamma values (Extended Data Table 1) and selected the best-performing model per organ. Dropout prevents overfitting by randomly disabling neurons during training, while gamma controls the learning rate decay.

#### Model selection criteria

To select the optimal model, we compared accuracy and loss using the following metrics:

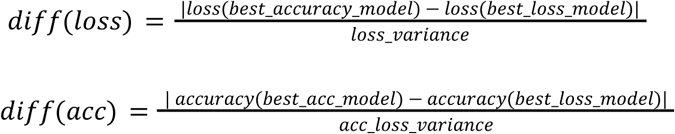

The final model was chosen based on the highest ratio, ensuring that the selected model provided the best trade-off between accuracy and loss improvement.

### Explainability analysis of CNN aging predictions using LIME

To interpret CNN aging predictions across different organs, we applied the Local Interpretable Model-Agnostic Explanations (LIME) method^43,93^. LIME is an explainability tool that generates local interpretations of classification models by perturbing input images and identifying regions that contribute to the model’s decision. We selected CNN classification predictions on the tiles in the validation set for both young and old classes. For each input tile, LIME produces a mask indicating positively predictive regions (green) that reinforce the classification and negatively predictive regions (red) that drive the decision in the opposite direction. This allows us to infer the image regions contributing to CNN aging predictions.

### Random Forests on gene expression TPMs for age classification

To assess organ-specific aging trajectories in gene expression for the ovary, uterus, and vagina, we trained separate Random Forest classifiers^94^ to distinguish young (<35 years) from old (>60 years) samples based on transcript-per-million (TPM) gene expression values.

For each organ, we identified the 1,000 genes with the most significant age-related expression changes using *limma*^95^. Raw TPM values of these genes were used as input features for model training. Hyperparameters were optimized using 5-fold cross-validation, and the best-performing model for each organ was selected based on classification accuracy.

### Vision Transformer for image segmentation of organs into tissues

To segment whole-slide images (WSIs) of reproductive organs into their constituent tissues and structures, we used the pre-trained ViT-S model, DINO ViT-S/8, following the pipeline described in Cisternino et al.^21^. Since the original model was not trained on all female reproductive organs (only in the breast), we performed two key adaptations: (1) fine-tuning the model on previously unrepresented WSIs of the reproductive organs and (2) introducing tissue-specific labels for each organ.

#### Model fine-tuning

We fine-tuned the model on WSIs from the uterus, ovary, vagina, ectocervix, endocervix, and fallopian tubes. First, we masked the 881 WSIs to separate tissue from the background using PathProfiler^96^. The foreground was then divided into 128×128 pixel tiles, yielding from an average of 62,000 tiles per whole slide image in the fallopian tubes up to 270,000 in the vagina (Extended Data Table Table 13). For fine-tuning, we randomly sampled 1,000 tiles per image and trained the model from the pretrained checkpoint^21^ for 50 epochs (until convergence, Extended Data Fig. 8A). The fine-tuned DINO ViT-S/8 was then used to generate 384-dimensional feature representations for all tiles.

#### Tissue labeling and segmentation

To classify tiles into specific tissue types, we first annotated representative tissue structures with the assistance of a histopathologist. Given the model’s high accuracy in generalization ability through clustering^21^ (Extended Data Fig. 2A), only a small number of images per organ needed manual annotation. We selected two WSIs per age group (young and old) based on GTEx pathological annotations, prioritizing inclusion of common histological categories (e.g., atrophy, inflammation). The histopathologist confirmed these annotations and outlined tissue regions. Using the annotations, we extracted coordinates of the labeled regions using MATLAB, mapping the outlined tissue regions to image tiles. Finally, we propagated labels to the remaining non-annotated tiles by computing feature distances and assigning labels using a k-Nearest Neighbors (k=200) algorithm.

### Genetic ancestry annotation

We inferred donor ancestry using genotype data from the GTEx v8 release^34^ by estimating ancestry proportions with ADMIXTURE (v1.3.0)^97^ To determine the optimal number of ancestry components (K), we tested models with two, three, and four populations. We selected K = 2 since it had the lowest cross-validation error. As a result, we obtained a continuous variable spanning from 0 corresponding to individuals of African ancestry to 1 corresponding to the European ancestry.

### Variance partitioning of ViT-derived image features

To identify key drivers of variation in image texture, we applied variance partitioning using a linear mixed model framework, implemented in the R package *variancePartition*^98^. This approach quantifies the proportion of variance in each image feature that can be attributed to fixed effects versus residual variance. For each tissue or tissue structure, we sampled 1,000 tiles per donor when available (e.g., excluding endometrium in myometrium-only samples or taking maximum available where <1000 tiles were present, Extended Data Table 16). For each donor-tissue pair, we computed the mean values of ViT-extracted image features and used these as input, along with the demographic and technical covariates, to fit the variance partitioning model. Each feature was modeled as:

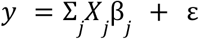

where *y* represents the image feature, *X* is the matrix of fixed effects, β − the coefficients for the fixed effects and ε - the residual variance.

We included continuous demographic traits and technical covariates commonly used in GTEx studies^99^, specifically, age, Body Mass Index (BMI), Genetic Ancestry (inferred from whole-genome sequencing^97^), Hardy Scale (donor death parameters) and Ischemic Time.

The final model formula used was:

*feature ∼ Age + BMI + Ancestry + HardyScale + IschemicTime*

By calculating the total variance, we determined the relative contribution of each fixed effect using the formula:

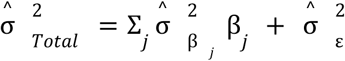

This allowed us to quantify the proportion of variation in image features explained by biological and technical factors.

### Visual Comparison of Morphological Differences by Age, BMI, and Genetic Ancestry

To examine morphology differences driven by age, BMI, and genetic ancestry, we selected representative donors guided by feature-level analysis and included additional examples in Extended Data Fig. 2.

For age-related comparisons, we selected three young and three old donors (or three middle-aged donors for vaginal epithelium and ectocervix) with the lowest and highest values, respectively, for the feature most strongly associated with age. In epithelial tissues, we examined an age-associated feature across the vaginal, endocervical, ectocervical, and fallopian tube epithelium, using the feature most strongly associated with age in the vagina. For cross-ancestry comparison, we selected six donors aged >50 (three of African and three of European ancestry) with low and high values, respectively, for the feature most strongly associated with ancestry in vaginal epithelium. For the BMI-related comparison, we selected six donors (three with BMI <23 and three with BMI >27) with low and high values, respectively, for the feature most strongly associated with BMI in breast adipose.

### Measurement of vaginal epithelium thickness using CellProfiler

To quantify vaginal epithelium thickness, we developed a CellProfiler pipeline^49^. As the input to the pipeline, a binary mask was generated for each whole-slide image by selecting all tiles classified as epithelium from the ViT-based segmentation output. Within the mask we identified continuous, elongated epithelial structures by tuning the *IdentifyPrimaryObjects* module to exclude small fragments and applying *FilterObjects* (eccentricity >0.85) to retain elongated regions. Thickness was measured using *MinorAxisLength* from the *MeasureObjectSizeShape* module, which fits an ellipse to each object and records its minor axis.

### Cell-type proportion changes with age

To further explore tissue composition changes with aging, we performed cell type deconvolution using the expression data for ovary and uterus and based on 2 cell atlases^100,101^ and compared the cell type proportion changes using cellular compositional data analysis (CoDA).

To estimate cell-type proportions in each organ, we applied GTM-decon^102,103^ to raw gene expression counts. This guided topic modeling approach deconvolves bulk transcriptomes by inferring cell-type-specific gene distributions from single-cell RNA-seq data. As this method requires a scRNA-Seq reference dataset for the inference, we used the Tabula Sapiens^100^ for the uterus, corresponding to one donor of 38 years old. For the ovary, we used a snRNA-Seq dataset for 8 donors aged from 23 to 54 years from a study of ovarian aging^100,101^. To measure the accuracy of the deconvolution, we split the single-cell data into training and test sets (in proportion 80:20), trained the model with the training set, and then simulated bulk data with the test set (we summed up all the counts per gene in all the cells as if we had one donor) and deconvoluted this “bulk” test set with the already trained model. With the ground truth cell-type proportions in the test set and the proportions obtained with the deconvolution, we calculated the Spearman R coefficient and the RMSE and compared them with the values obtained in the description of the method. For each organ, we trained the models for 5, 25, and 50 iterations and selected 5 iterations as optimal based on accuracy (Extended Data Table 18).

To investigate age-related changes in cell-type proportions, we performed a cellular Compositional Data Analysis (CoDA). First, we transformed relative cell-type proportions into pivot coordinates^104,105^ as in Cisternino et al.^21^. We then applied a linear mixed model to assess how cell-type proportions change with age while accounting for demographic variables (BMI, genetic ancestry) and technical covariates (HardyScale, IschemicTime, RIN, ExonicRate) (Supplementary Formula 1). Multiple testing correction was performed using the Benjamini–Hochberg false discovery rate (FDR), and cell types were considered to have significant age-associated proportional changes at FDR-adjusted p < 0.05.

**Supplementary Formula 1**

*freq ∼ HardyScale + IschemicTime + RIN + ExonicRate + Age + Ancestry + BMI*

### Tissue-specific aging classification with Elastic Net

To classify aging patterns within specific tissue structures, we trained Elastic Net classifiers using self-learned image representations extracted from the fine-tuned ViT-S/8 model.

For each models’ training, we combined the training and testing sets used for CNN training, again using only young (<35 years) and old (>60 years) samples. For each tissue or tissue structure, we sampled 1,000 tiles per donor when available (e.g., excluding endometrium in myometrium-only samples or taking maximum available where <1000 tiles were present, see Supplementary Note). Each tile’s 384-dimensional feature representation was extracted using the fine-tuned ViT-S/8 model.

Elastic Net classifiers were trained to distinguish young from old tiles using these features. Model hyperparameters were optimized via 10-fold cross-validation, ensuring that tiles from the same donor were kept within either the training or test sets. The final model was validated on the separate validation set to assess classification accuracy, and all tiles were classified to infer aging trajectories.

### Gene quantification

Gene quantification was based on GENCODE v39 annotations^106^ (https://www.gencodegenes.org/human/release_39.html). We obtained gene counts and TPM values from the GTEx portal (https://gtexportal.org/home/datasets) and filtered for protein-coding and lncRNA genes. For expression analysis, we included genes meeting these criteria: TPM ≥ 1, counts ≥ 10, and expression in ≥ 20% of tissue samples, excluding genes in pseudoautosomal regions. In total, we analyzed 21266 genes (15914 protein-coding and 5352 lincRNA) across organs.

### Analysis of image- and gene expression-based aging trajectories

To assess aging trajectories across image- and gene expression-based predictions, we estimated the probability of being classified as old for all samples using three different types of trained classifiers. For images, we used organ-wide CNNs on large histological tiles and tissue-specific Elastic Net classifiers on ViT-derived tile features. For gene expression, we used Random Forest models on bulk gene expression TPMs for each organ. These probabilities formed aging trajectories, representing the distribution of “old” classification probabilities across age.

To determine if an aging trajectory followed a linear pattern, we applied the Davies test for non-zero slope differences^102^ using the *davies.test* function from the R package *segmented*. If a trajectory exhibited a significant slope change (p < 0.05), we identified the age of the most rapid transition.

We first smoothed the trajectory using LOESS (Locally Estimated Scatterplot Smoothing) with a span of 0.75 and generated high-resolution age predictions across the range 20 to 70 years (200 evenly spaced points). The age with the most rapid change was identified as the timepoint with the largest slope, calculated as the first derivative of the smoothed probability function:

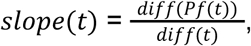

where *t* is age and *Pf*(*t*) is the LOESS-smoothed probability of classification as old.

### Integration of image features and gene expression using Multi-Omics Factor Analysis (MOFA)

To jointly analyze image features and gene expression data and explore their correlation with covariates, we applied Multi-Omics Factor Analysis (MOFA)^50,102^. MOFA identifies latent factors that explain variability across multiple data modalities measured on the same set of donors, allowing us to correlate these factors with metadata variables.

We extracted 384 image features per tile using the fine-tuned ViT-S/8 model, then averaged these features per tissue per donor, creating a dataframe with 384 rows (features) and X columns (donors) per tissue. As a second data modality, we included gene expression data, selecting the 5,000 most variable genes based on normalized expression counts, as recommended in MOFA guidelines.

Before running MOFA, we adjusted for technical covariates for both data modalities by fitting a linear model to each image feature and running voom-limma pipele on gene expression data, and taking into the MOFA the corresponding residuals. For the images and gene expression we had different available technical covariates:

- Image features: Ischemic Time, Hardy Scale.
- Gene expression: Ischemic Time, Hardy Scale, PEER2, NucIsoBatch, RIN, and Exonic Rate.

Following MOFA inference, we examined the correlation between the extracted factors and age. Additionally, we performed Gene Set Enrichment Analysis (*clusterProfiler*) with Gene Ontology database (GO) for Biological Processes on genes ranked by their weight within factors shared between gene expression and image features that were significantly correlated with age.

### Differential Gene Expression Analysis

To identify the differentially expressed with age genes (age-DEGs), we used linear regression models following the voom-limma pipeline^95,107^. Excluding the organs with sample size <20 (endocervix, ectocervix and fallopian tubes), we conducted the analyses separately for each of the four organs and the myometrium. We included samples (n = 659) from donors (n = 245) with complete metadata for technical covariates routinely used in previous GTEx publications^99,108^ and demographic traits including genetic ancestry, age, and body mass index (BMI). The technical covariates accounted for parameters related to donor death, ischemic time, RNA integrity number (RIN), and sequencing quality metrics. To control for unmeasured sources of variation, we corrected for PEER factors 1 and 2, which primarily capture cell type heterogeneity^109^ and sequencing batch effects^110^, respectively. Correlation analysis between PEER factors and tissue proportions (Extended Data Fig. 8B) indicated low correlation values, allowing us to retain PEER1 and PEER2 factors to preserve biological information.

To identify age-DEGs independently of tissue composition, we incorporated the proportions of tissues in each organ extracted from the ViT-based image segmentation (Extended Data Tables 7,9,11-12). We converted the proportion values of each tissue to pivot coordinates as in Cisternino et al.^21^ to account for the dependence of compositional values^104^:

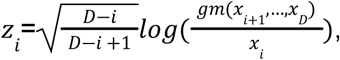

Where:

*D* is the number of components (proportions), *i* is the index of the current component, *gm*(*x*_*i*+1_,…,*x*_*D*_) is the geometric mean of the components from *i+1* to *D*, and *x*_*i*_ is the value of the *i*-th component.

We applied multiple testing correction to all analyses using the Benjamini–Hochberg false discovery rate (FDR) method and classified genes as differentially expressed if they had an adjusted p-value below 0.05.

**Formula 1**

*Expression (log2cpm) ∼ HardyScale + IschemicTime + RIN + Cohort + NucIsoBatch +ExonicRate + PEER1 + PEER2 + Age + Ancestry +BMI + tissue proportions*

### Addressing Uterus Sampling Challenges

A major challenge in uterus analysis was inconsistent sampling, as 43% of samples lacked endometrium (Extended Data Fig. 4C). Endometrium proportion varied widely within age groups (Extended Data Fig. 4D), likely due to sampling and non-annotated biological effects, such as monthly remodeling due to menstrual cycle in young women^111^, and known differences in postmenopausal atrophy^112^. To address this, we performed all the described analyses where we analyzed endometrial properties (classification, variance partition and MOFA) only in samples containing both endometrium and myometrium.

To determine whether myometrium’s higher variance was biologically driven rather than a sampling artifact, we repeated variance partitioning on samples containing both tissues, confirming that myometrium consistently showed greater age-related variance (Extended Data Fig. 8C).

Additionally, differential expression analysis of myometrium-only samples, correcting for vessel proportions, yielded a total of 1159 age-DEGs specific to myometrium (Fig. 4A).

### Variance partitioning and tissue classification on downsampled data

Uneven tissue representation (Extended Data Fig. 4B) could bias comparisons when certain tissues are underrepresented. However, even when a sample is dominated by one tissue type, our tile-based approach allows us to extract tens to hundreds of tiles from the minority tissue (Extended Data Fig. 8D, Extended Data Table 16). Across organs, we found no association between tile count and age-related variance (Extended Data Fig. 8D). To further validate this, we performed downsampling specifically in the uterus—where the largest tissue imbalance occurs—and confirmed that myometrium consistently exhibited the highest age-related variance, with aging trajectories and classification accuracies remaining unaffected (Extended Data Fig. 8E-G).

### Sliding-window Differential Expression Analysis

To investigate age-associated changes in gene expression across the age range (21-70), we conducted a sliding-window differential expression analysis. This approach follows the differential expression analysis framework described above, but applies a moving window across the age range.

For each window center (w) spanning ages 21 to 70, sliding by 1 year, we defined a bucket size (b) of 10 years. In each iteration, samples were assigned to one of two age bins:

- **Age bin = 0:** Samples with ages in the interval [w−b,w).
- **Age bin = 1:** Samples with ages in the interval (w,w+b].

Differential expression analysis (DEA) was then performed using the following linear model:

*Expression (log2cpm) ∼ HardyScale + IschemicTime + RIN + Cohort + NucIsoBatch +ExonicRate + PEER1 + PEER2 + Age bin + Ancestry + BMI*

Multiple testing correction was applied using the Benjamini–Hochberg FDR, with genes classified as differentially expressed at an adjusted p-value < 0.05. This approach allowed us to assign differentially expressed genes per window center.

### Clustering of common aging-expression trajectories

To identify common aging-related gene expression trajectories, we applied hierarchical clustering based on Euclidean distances between loess-smoothed trajectories (span = 0.75) of z-scored gene expression counts^65^. For clustering, we used *hclust* function with *complete* clustering method from R package *cluster*. The input genes were those identified as differentially expressed with age in the differential expression analysis (DEA) (Formula 1).

The number of clusters was determined proportionally to the size of the differentially expressed gene sets, resulting in 10 clusters for the ovary, 14 for the uterus, and 2 for the vagina.

### Functional Enrichment

To understand the potential functional significance of the genes of each cluster defined in the clustering analysis, we conducted an OverRepresentation Analysis (ORA) using GeneOntology database, with *clusterProfiler* package^65,113^. We accounted for multiple testing using the Benjamin-Hochberg method, with FDR <0.05. To ensure context-specific analysis, our Gene Universe for background for each organ was all the genes expressed in the organ.

### Enrichment of age-DEGs with GWAS traits

GWAS traits included largest studies for reproductive aging traits (age at menarche and age at menopause)^62,114^, common clinical conditions (pelvic organ prolapse, polycystic ovary syndrome, endometriosis, polyps of the female genital tract, uterine fibroids and ovarian cysts)^72,115–119^ and hormonal traits (sex-hormone binding globulin, estradiol, bioavailable testosterone and anti-Müllerian hormone levels)^120,121^ (Extended Data Table 13). Candidate gene lists for those traits were sourced either from the GWAS Catalog^122^ or manually curated from each respective study’s presented data (Extended Data Table 13). Then, per each organ or tissue, we assessed whether age-DEGs were enriched for these trait-associated genes using the fisher.test function from R (v4.1.2) with default parameters. To ensure context-specific analysis, the list of background genes for each organ was all the genes expressed in the organ. We applied the Benjamini–Hochberg method to adjust for FDR, considering a tissue to be significantly enriched with age in a particular gene set at FDR < 0.05.

### Segmentation of ovarian follicles

It is important to note that we were unable to segment follicles in the ovary—a small, fluid-filled sac in the ovary that contains one immature egg and key functional unit responsible for an individual’s reproductive potential^111,123^. This limitation arises from the significant variation in follicle morphology and size across developmental stages (primordial, primary, preantral, secondary, tertiary)^123^, which would require a specialized classifier for accurate annotation. Additionally, large antral follicles, characterized by their fluid-filled cavities, often resembled cysts or tissue tears, leading to misclassification or exclusion.

### LLM usage

AI-based tools were used to streamline and edit the manuscript text. All content and interpretations are the sole responsibility of the authors.

## CODE AVAILABILITY

### Data and code availability

The data used for the analyses described in this manuscript were obtained from: the [GTEx Portal](https://www.gtexportal.org/home/) v10 (counts and TPMs) on 11/27/2024 and [dbGaP](http://www.ncbi.nlm.nih.gov/gap) accession number [phs000424.v8](https://www.ncbi.nlm.nih.gov/projects/gap/cgi-bin/study.cgi?study_id=phs000424.v10.p2) (metadata).

Analysis scripts and pipeline are available at github: https://github.com/Mele-Lab/2025_GTEx_Menopause

## ACKNOWLEDGEMENTS

N.P.G. acknowledges her AI4S fellowship within the “Generación D” initiative by Red.es, Ministerio para la Transformación Digital y de la Función Pública, for talent attraction (C005/24-ED CV1). Funded by the NextGenerationEU funds through PRTR. M.M. was supported by a grant PID2019-107937GA-I00 funded by MCIN/AEI/10.13039/501100011033 and a grant RYC-2017-22249 funded by MCIN/AEI/10.13039/501100011033 and by “ESF Investing in your future”.

We are deeply grateful to Winona Oliveros for her invaluable input and timely advice, providing the genetic ancestry annotation. We thank Camil Castelo-Branco for his insightful gynecological perspective on reproductive aging. Lastly, we are grateful to the Melé laboratory at the Barcelona Supercomputing Center (BSC) for their support, discussions, and valuable feedback throughout this project.

Figure 1A was done with BioRender.com.

## SUPPLEMENTARY FILES

**Extended Data table 1.** Parameters of training, accuracy, and sample sizes in training, testing and validation of the age group CNN classifiers.

**Extended Data Table 2.** Tests for nonlinearity and changepoints. Davies test performed on model predictions across age. Age of change defined as age of maximum slope (see methods).

**Extended Data Table 3.** Random forest classifiers statistics across organs.

**Extended Data Table 4.** MOFA factors’ correlation with age and their corresponding adjusted p-values (left) and variance explained of each data modality per factor within each organ.

**Extended Data Table 5.** GSEA with GO terms of Biological Processes (BP) of the ranked 5000 HVGs according to their weight for factors significantly correlated with donor age (FDR<0.05).

**Extended Data Figure 6.** Sample sizes and accuracies for tissue-specific Elastic Net classiffiers.

**Extended Data Table 7.** Differentially expressed with age genes in uterus and their cluster attribution.

**Extended Data Table 8.** Uterus. Enrichments for clusters. Functional Enrichment with Gene Ontology for gene clustered according to their aging dynamics (Fig 4 C-E, Ext Data Fig 5-6). Background is all genes expressed in the organ.

**Extended Data Table 9.** Differentially expressed with age genes in ovary and their cluster attribution.

**Extended Data Table 10.** Ovary. Enrichments for clusters. Functional Enrichment with Gene Ontology for gene clustered according to their aging dynamics (Fig 4 C-E, Ext Data Fig 5-6). Background is all genes expressed in the organ.

**Extended Data Table 11.** Differentially expressed with age genes in vagina and their cluster attribution + Enrichment with GO:BP for the significantly enriched cluster.

**Extended Data Table 12.** Differentially expressed with age genes in breast.

**Extended Data Table 13.** Reproductive studies description: sources of data for the targeted selection for the reproductive GWAS included.

**Extended Data Table 14.** Results for the GWAS enrichment (Fisher’s test) between DEGs with age in each reproductive organ or tissue and the selected GWAS studies (tissues and traits with at least 1 overlapping gene are shown).

**Extended Data Table 15**. Image filtering terms. For each organ we removed from the analyses the mislabeled images based on GTEx histopatholgical annotations.

**Extended Data Table 16.** Number of tiles per organ and per tissue: Total available and numbers that entered into analyses.

**Extended Data Table 17.** Davies’ test for proportions of each tissue of each organ across ages, to test the non-linearity of the trajectories.

**Extended Data Table 18.** Deconvolution accuracy measured with Spearman R coefficient an root-mean-square error (RMSE) between the simulated bulk data from the test set and the deconvoluted single-cell data from the training set.

## SUPPLEMENTARY FIGURES

**Extended Data Fig. 1.**
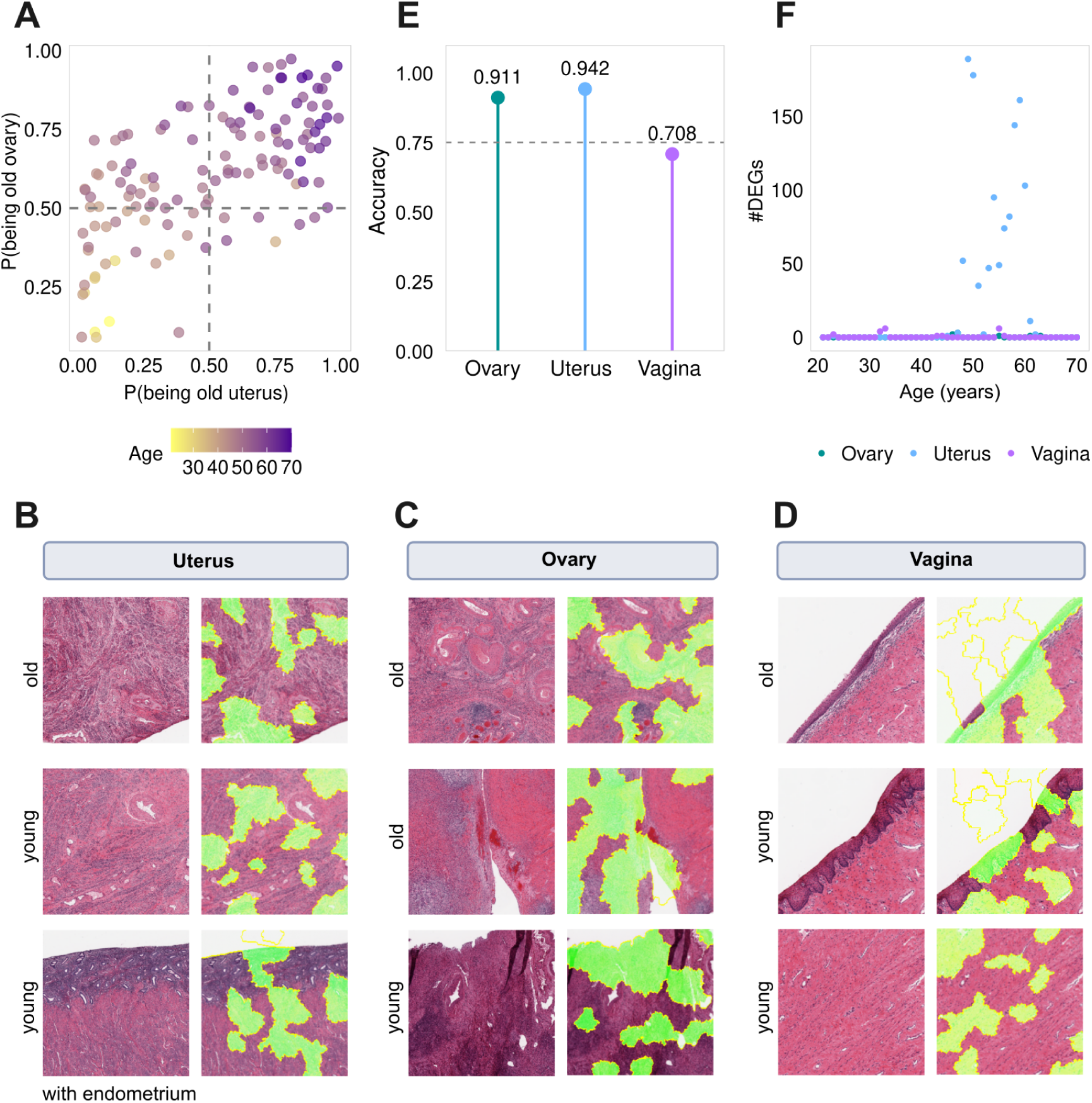
Age classification and visual interpretation from histology and gene expression across female reproductive organs. A CNN-predicted probabilities of being classified as “old” based on ovary or uterus images from the same donor. B-D Representative histological tiles from young and old samples for the uterus (B), ovary (C), and vagina (D) (left); corresponding LIME explanations highlighting image regions (green) that increase CNN model confidence (right). E Classification accuracies of random forest age-group classifiers using gene expression TPMs. F Number of differentially expressed genes with age in the uterus, ovary, and vagina across 20-year sliding windows (1-year offset).

**Extended Data Fig. 2:**
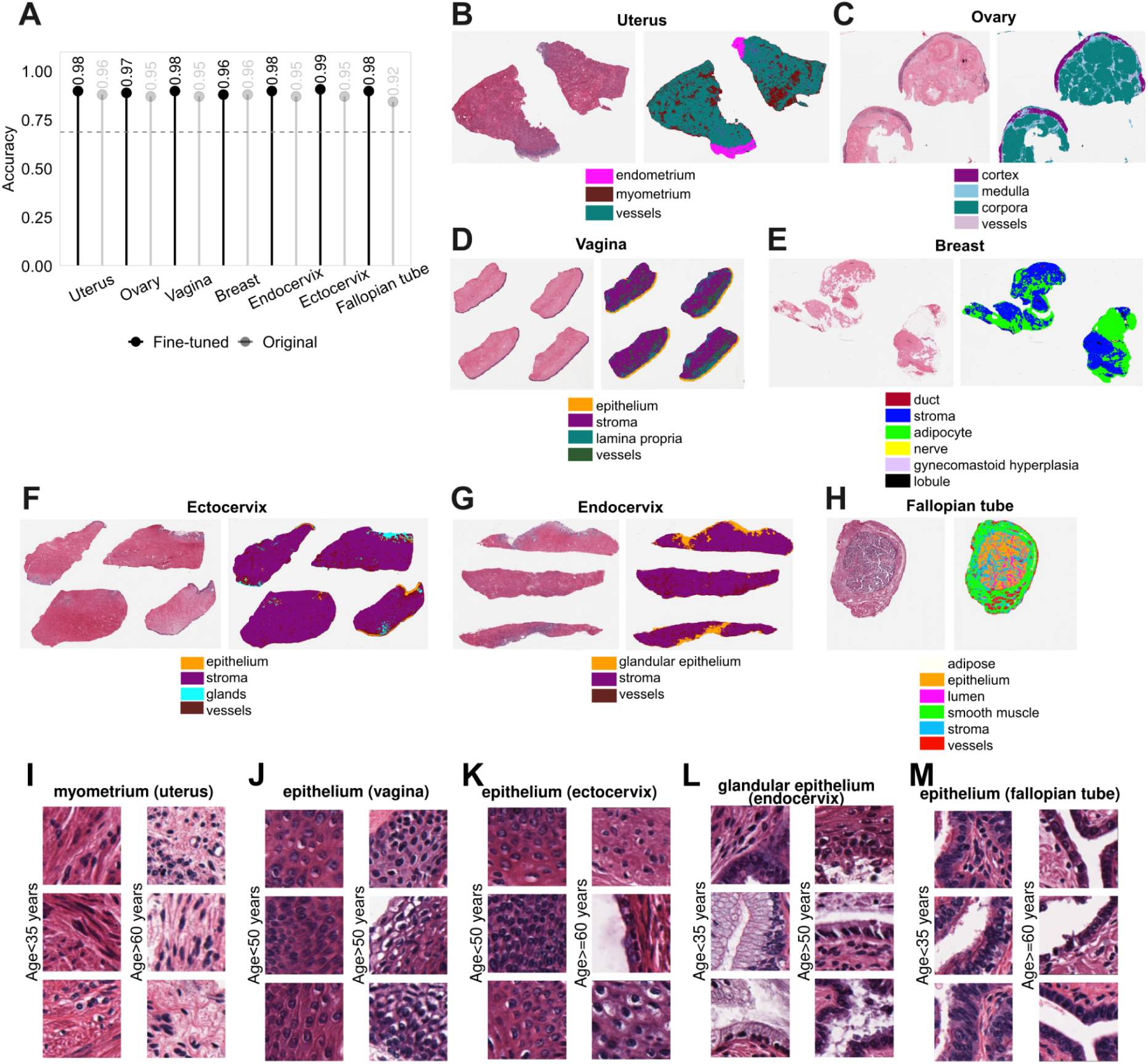
Vision transformer histology organ segmentation and tile examples. **A** Clustering accuracies per donor and tile across organs. B-H Organ segmentation examples showing tissue and structure classifications. **I-M** Age-related histological changes in myometrium (I), and a higher nuclear-to-cytoplasm ratio in epithelia of vagina (J), ectocervix (K), accompanied with the flattening of columnar cells endocervical glandular (L), and fallopian tube epithelium (M) in old donors^48^.

**Extended Data Fig. 3.**
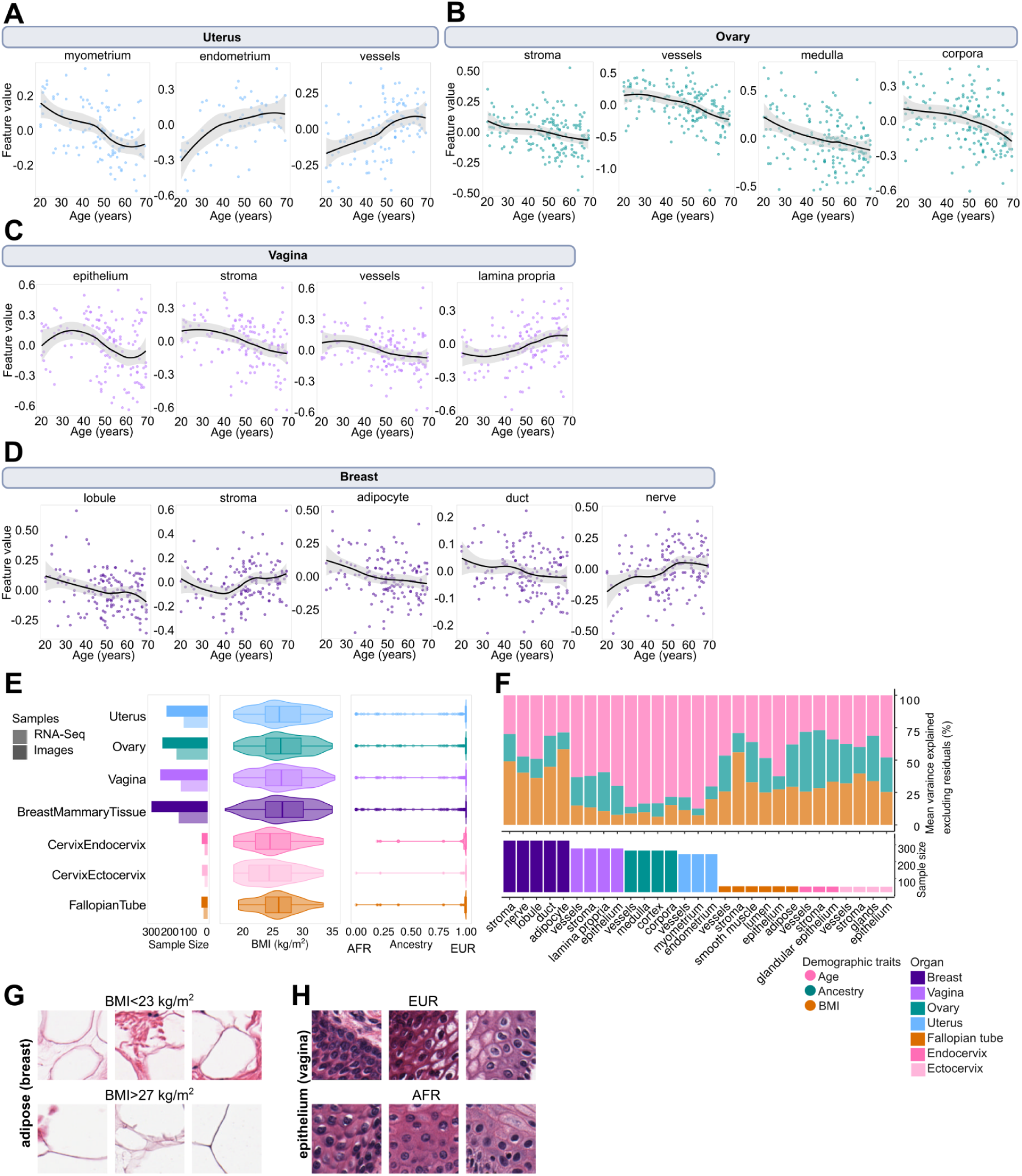
Image features trajectories and demographic variability across donors and organs. **A-D** Feature image trajectories for each tissue structure in uterus (A), ovary (B), vagina (C), breast (D). **E** BMI and genetic ancestry distributions of image samples. **F** Mean variance explained by demographic traits across tissues.**G** BMI-related histological differences in breast adipose tissue, with larger adipocytes and reduced intercellular space in high-BMI donors. **H** Genetic ancestry-associated differences in vaginal epithelium, appearing denser in donors of African ancestry compared to European ancestry.

**Extended Data Fig. 4.**
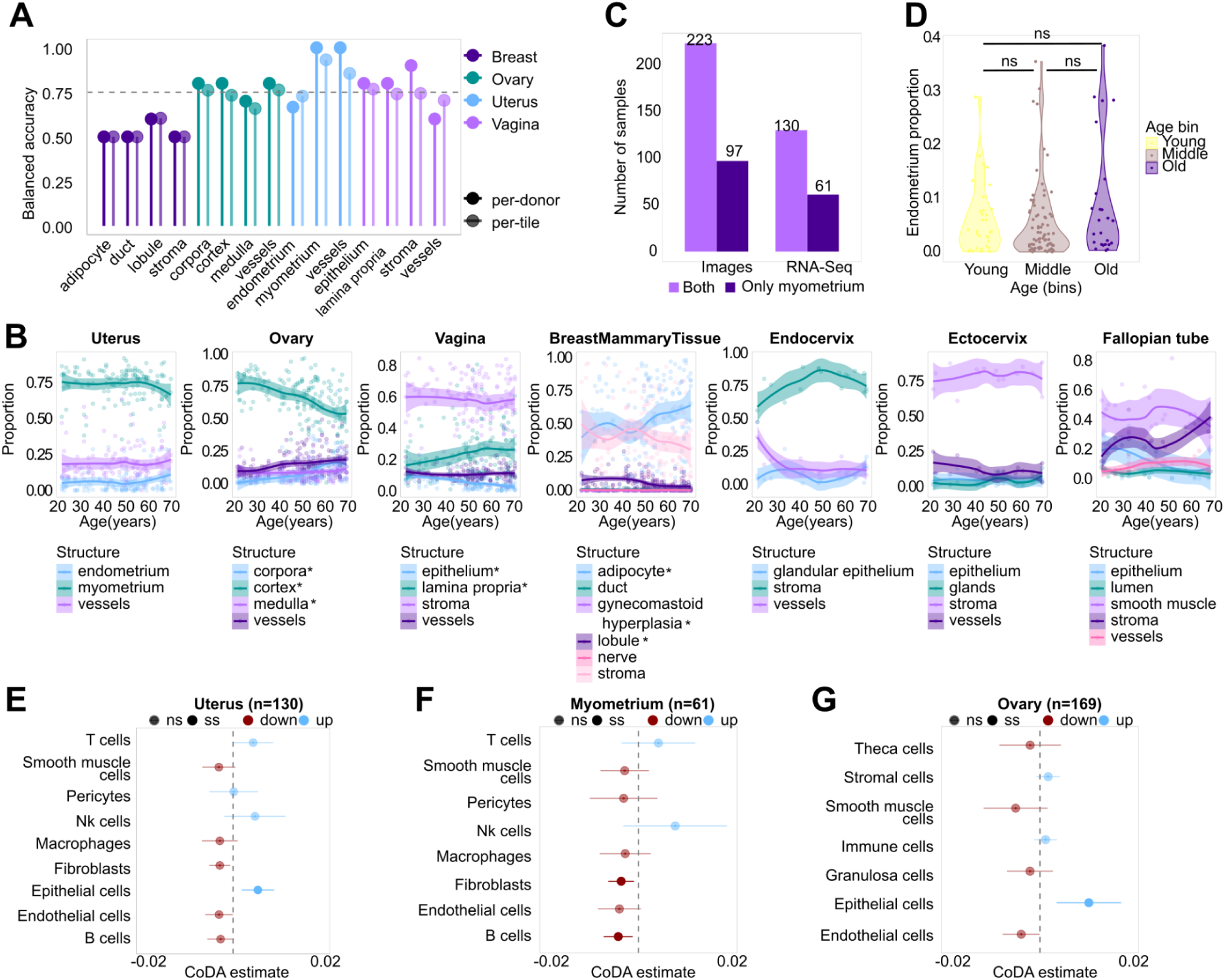
Tissue-specific and cell type-specific aging dynamics. **A** Balanced accuracies for tissue-specific Elastic-Net classifiers. **B** Tissue and tissue structure proportions across organs. Asterisk (*) denotes the tissues that change significantly with age (FDR<0.05) in the compositional data analysis. **C** Number of images and RNAseq samples with and without endometrium. **D** Proportion of endometrium within uterus samples across age groups. **E-G** Cell-type proportion changes with age for samples of uterus (E), myometrium (F) and ovary (G).

**Extended Data Fig. 5.**
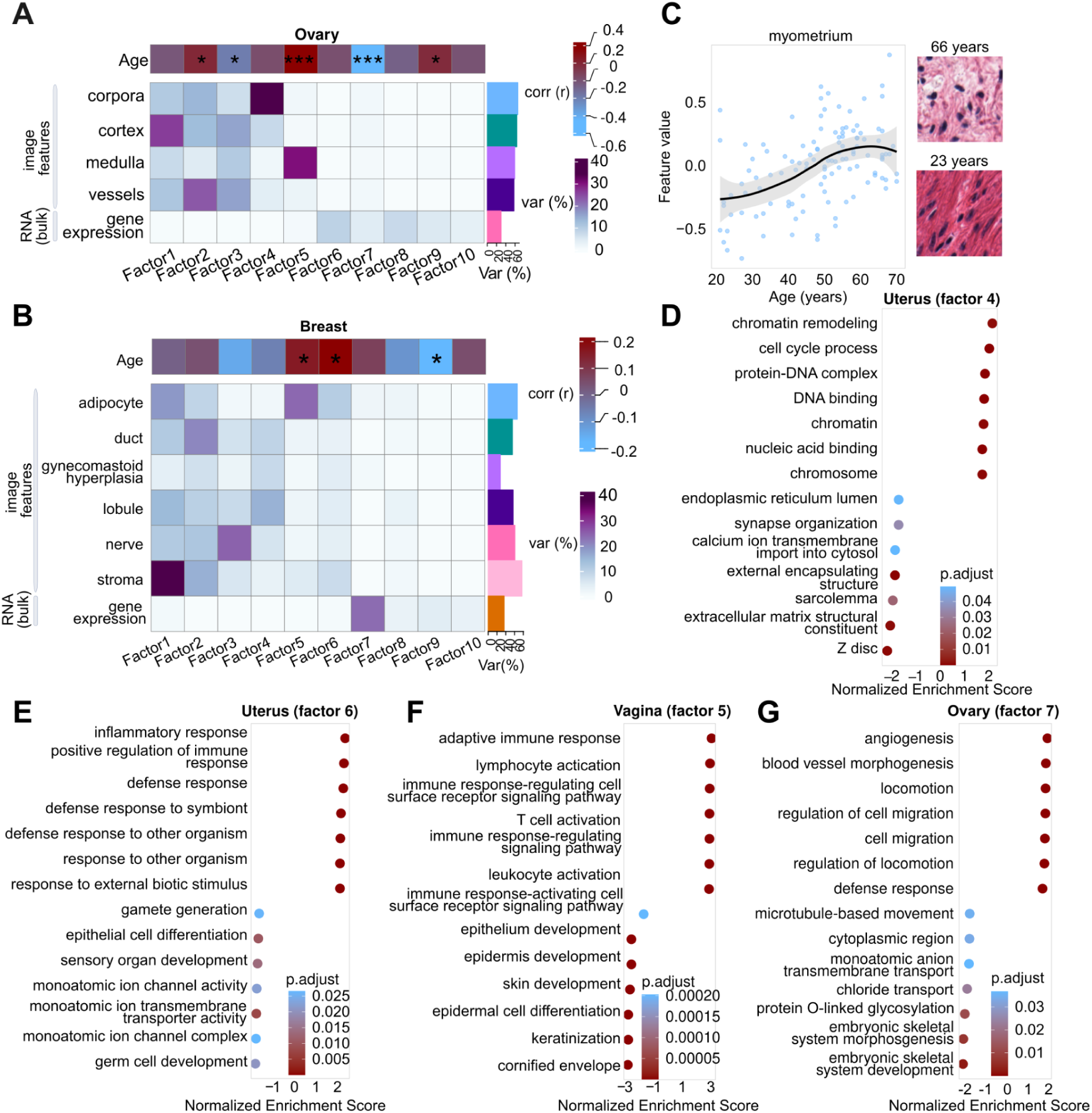
Multimodal analysis for ovary and breast and up and downregulated functions with age. **A-B** Multi-Omics Factor Analysis (MOFA) results for ovary (A) and breast (B): heatmaps show variance explained by 10 latent factors across modalities (bottom) and their correlation with donor age (top). Asterisks denote significance (*FDR<0.05 & >0.005, **FDR<0.005 & >0.0005, ***FDR<0.0005). **C** Trajectory of the most weighted feature for Factor 4 in myometrium, showing the difference between low values in young donors (left) and high values in old donors (right). **D-G** Gene Set Enrichment Analysis (GSEA) of the top 7 genes with positive (NES>0) and negative (NES<0) normalized enrichment scores using Gene Ontology: genes contributing to Factor 4 in the uterus (D), Factor 6 in the uterus (E), Factor 5 in the vagina (F), and Factor 7 in the ovary (G).

**Extended Data Fig. 6.**
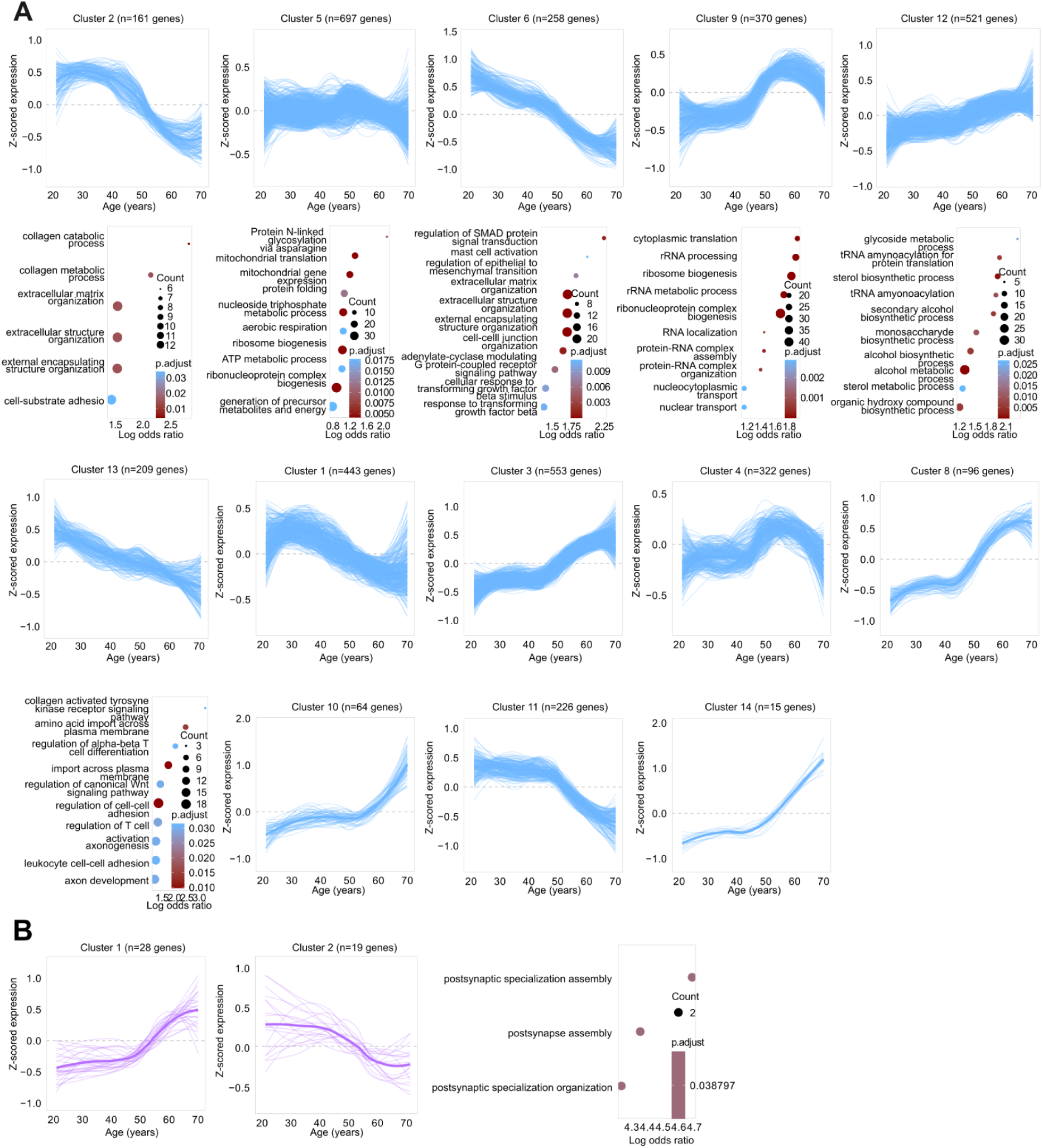
Gene expression trajectory clustering across organs. **A** Gene count aging trajectories in the uterus, with Gene Ontology enrichments shown below clusters with significant enrichment. **B** Gene count aging trajectories in the vagina, with Gene Ontology enrichment displayed next to the significantly enriched cluster.

**Extended Data Fig. 7.**
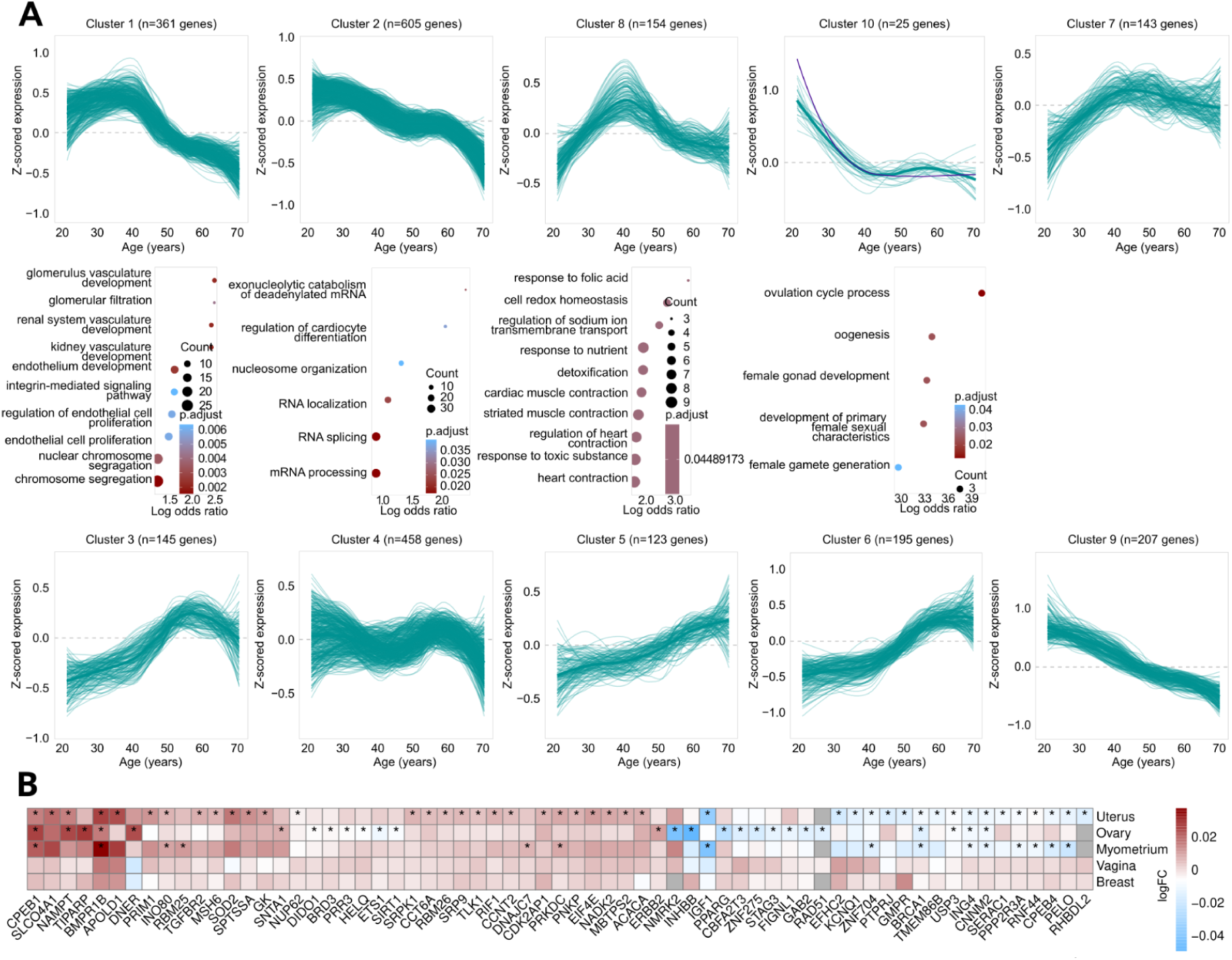
Gene expression trajectory clustering across organs and GWAS overlap. **A** Gene count aging trajectories in the ovary, with Gene Ontology enrichments shown below significantly enriched clusters. **B** Heatmap for 65 age-DEGs in the uterus, ovary, and/or the myometrium overlapping with menopause-associated GWAS genes. Asterisks indicate significant age-DEGs (FDR<0.05). Grey squares indicate missing data.

**Extended Data Fig. 8.**
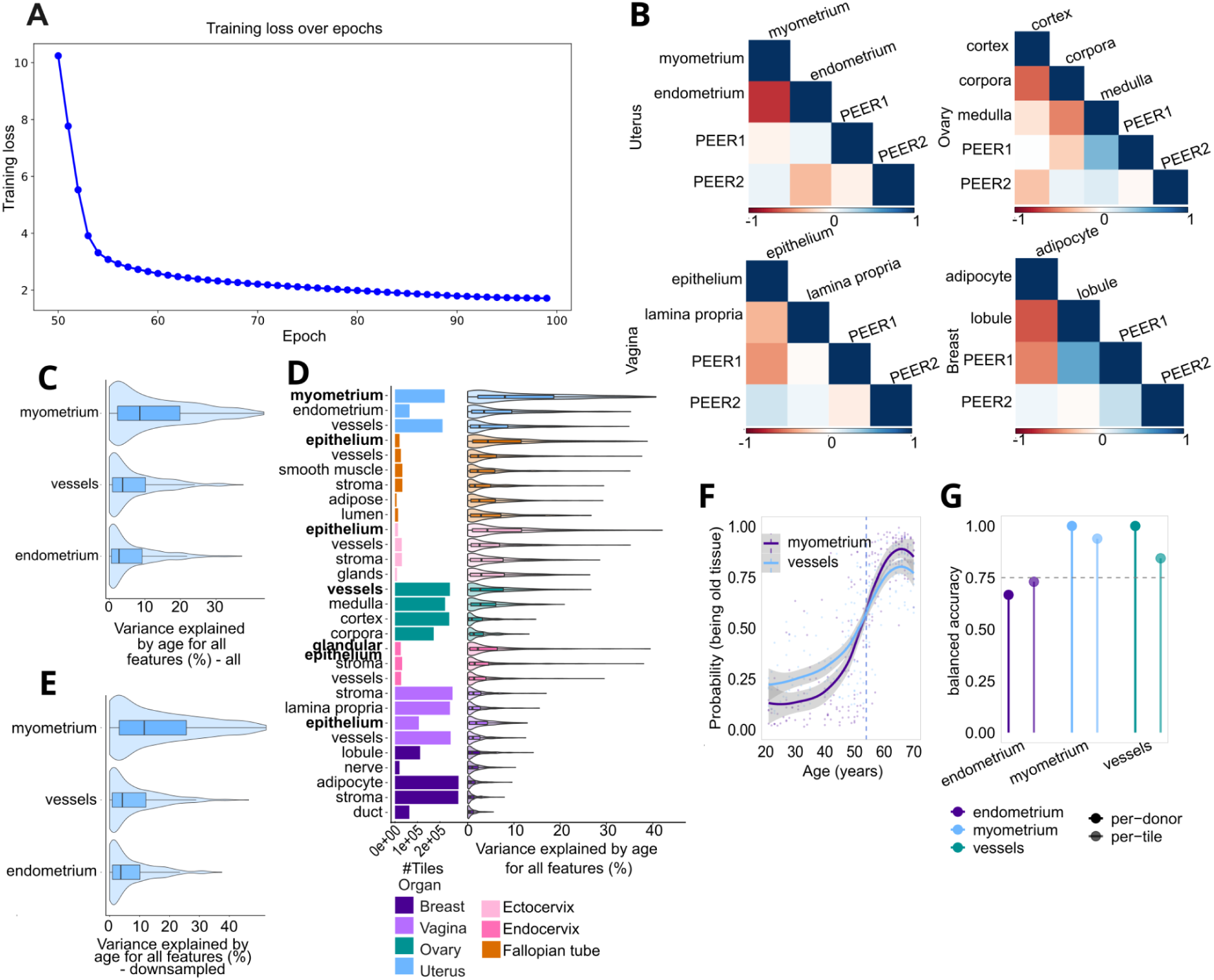
Methods. **A** Curve of training loss across epochs. **B** Correlation values between PEER factors and tissue proportions. **C** Variance explained by age in uterine tissues in the samples that have both endometrium and myometrium. **D** Variance explained by age by all tissues with overall tile numbers per tissue (analyses were conducted on randomly sampled 1000 tiles per tissue). **E** Variance explained by age in uterine tissues in downsampling analysis. **F-G** Uterine tissue classification trajectories in downsampling analysis (F) and classification accuracies (G).

